# *Withania somnifera* root extract (LongeFera™) confers beneficial effects on health and lifespan of the model worm *Caenorhabditis elegans*

**DOI:** 10.1101/2024.08.01.606126

**Authors:** Nidhi Thakkar, Gemini Gajera, Dilip Mehta, Sujit Nair, Vijay Kothari

## Abstract

*Withania somnifera* is among the most widely prescribed medicinal plants in traditional Indian medicine. Hydroalcoholic extract of the roots of this plant was investigated for its effects on the overall health and lifespan of the model worm *Caenorhabditis elegans*. Extract-exposed worms registered longer lifespan, higher fertility, better motility and metabolic activity. Whole transcriptome analysis of the extract-treated worms revealed the differential expression of the genes associated with lifespan extension, eggshell assembly and integrity, progeny formation, yolk lipoproteins, collagen synthesis, cuticle molting, etc. This extract seems to exert its beneficial effect on *C. elegans* partly by triggering remodeling of the developmentally programmed apical extracellular matrix (aECM). Differential expression of certain important genes (*cpg-2*, *cpg-3, sqt- 1, dpy-4, dpy-13,* and *col-17*) was confirmed through PCR assay too. Some of the differently expressed genes (*gfat-2, unc-68, dpy-4, dpy-13, col-109, col-169,* and *rmd-1*) in worms experiencing pro-health effect of the extract were found through cooccurrence analysis to have their homologous counterpart in humans. Our results validate the potential of *W. somnifera* extract as a phytopharmaceutical.

## 1. Introduction

Since centuries, humans have longed for a long and healthy life. A variety of plant extracts and polyherbal formulations have been claimed in different systems of traditional medicine to impart the same. However widespread application of such plant products in modern times asks for scientific validation of the claimed activities. One of the most popular plants in Indian system of *Ayurved* is *Withania somnifera*. Commonly known as *Ashwagandha*, this plant belongs to the Solanaceae family, and has a long history of >3000 years of being prescribed in indigenous medicine (Bharti et al., 2016). In *Ayurved*, *W. somnifera* has been referred to as ‘Rasayan’, which means a holistic therapy for suppressing the aging process, developing positive physical and mental health, and boosting the immune system and maintaining youth. Ancient texts mention *W. somnifera* as ‘Avarada’ whose meaning is related to longevity or regeneration. The multitude of activities claimed in this plant include immunomodulatory (Singh et al., 2022), anti-inflammatory (Sharma et al., 2023), neuroprotection (Kuboyama et al., 2014), pro-fertility (Sengupta et al., 2018), anticancer (Mehta et al., 2021), and anti-stress (Speers et al., 2021). Its regular consumption is believed to rejuvenate the reproductive organs and promote fertility, retard senescence, relieve nervous exhaustion, sexual debility, muscular weakness and geriatric problems. *W. somnifera* is expected to inhibit aging and catalyze the anabolic processes of the body (Baliga et al., 2015). Among all parts of this plant, its root has received maximum attention in medicinal texts and practice. This is because its roots have the highest concentration of desired bioactive principles (Basudkar et al., 2024). Further, with respect to safety too, only the root part of *W. somnifera* is recommended for therapeutic and internal administration (Vaidya et al., 2024).

Mainstreaming the use of any herbal preparation for therapeutic as well as nutraceutical applications is dependent on demonstration of claimed biological activities in them through appropriate scientific assays. Owing to concerns of ethics and feasibility, assays with higher animals and human volunteers are often challenging. Lower animals like the nematode worm *Caenorhabditis elegans* offer a useful and relatively convenient and biologically relevant platform for assessment of biological activities in natural as well as synthetic preparations at the whole organism level. In recent years, wild type and transgenic strains of *C. elegans* have been employed widely as model organisms for assays relevant to neurology (Calahorro et al., 2011), lifespan (Amrit et al., 2014), diabetes (Zhu et al., 2016), wound healing (Xu et al., 2012), and microbial virulence (Sifri et al., 2005). Aging being a complex and multifactorial phenomenon, is difficult to be modelled, and more so the assessment of longevity or lifespan within a shorter time frame. Though *C. elegans* has a short lifespan (few days), fundamental biological mechanisms and systems in this worm are similar to the mammalian system (Arya et al., 2010). The mechanisms associated with increase in longevity identified in this worm are shown to follow a pattern similar to those in humans (Kojima et al., 2004; Suh et al., 2008). Though lifespan-enhancing effect of *W. somnifera* root extract has been reported earlier too (Kumar et al., 2013), much remains to be elucidated with respect to the underlying molecular mechanisms. This study attempted to fill the said gap by investigating effect of the hydroalcoholic extract of *W. somnifera* root on lifespan and overall healthspan of *C. elegans* through *in vivo* assays as well as gene expression analysis at the whole transcriptome level.

## 2. Methods

### 2.1. Plant extract

The hydroalcoholic extract (LongeFera^™^) of *Withania somnifera* root was procured from Phytoveda Pvt. Ltd., Mumbai. The roots of *W. somnifera* were collected from Madhya Pradesh, India. It was authenticated by a taxonomist at Botanical Survey of India, Jodhpur, and the voucher specimen was deposited there with reference ID: BSI/AZRC/I.12012/Tech/19-20/PI.Id/671. The root material was washed, dried, and then pulverized. The resulting powder was then extracted with ethanol: water (8:2 v/v) at 60 °C. Powdered form of this extract was then used for assay purpose. The extract was analysed as per United States Pharmacopeia (USP) for various quality control parameters (impurities, heavy metals, microbial load, etc.). The content of withanosides and withanolides was determined using HPLC. Major phytocompounds identified in the extract along with their concentration are depicted in the supplementary chromatogram (Figure S1). Total withanolides content in the extract was found to be 2.69 ± 0.02%.

The extract powder was suspended in water or DMSO (Merck) for bioassay purposes. While the extract was fully soluble in DMSO, its insoluble fraction from the aqueous suspension was removed by centrifugation (7500 g at 25°C) for 10 minutes. The soluble fraction (supernatant) was filtered through a syringe filter (0.45 µm; Axiva), and the filtrate was stored in a sterile 15 mL glass vial (Borosil) under refrigeration. The solubility of the extract in water was 81.84%.

### 2.2. Test organism

Wild type N2 Bristol strain of *Caenorhabditis elegans* procured from the Caenorhabditis Genetics Center (USA) was used as model organism in this study. Lyophilized *E. coli* OP50 procured from Biovirid (The Netherlands) was used as food for *C. elegans*, while maintaining the worm on NGM agar plates (Nematode Growing Medium; 3g/L NaCl, 1M CaCl2, 1M MgSO4, 2.5g/L peptone, 5 mg/mL cholesterol, 1 M phosphate buffer of pH 6, 17g/L agar-agar). Worm synchronization was done as described in literature (Corsi et al., 2015) and in our previous studies (Joshi et al., 2019; Patel et al., 2019) too. Prior to the *in vivo* assays, worms were kept without food for two days to make them gnotobiotic.

### 2.3. Lifespan assay

Synchronized (L3-L4 stage) gnotobiotic worms were incubated at 22±1°C with different concentrations (5-1000 µg/mL) of the plant extract in 24-well plates (HiMedia). Each well contained 1 mL total volume (995 µL M9 buffer + 5 µL extract). Control wells contained only M9 buffer with worms, but no extract. Ten worms were added in each well, and these 24-well plates were monitored over a 12-day period (till all worms died in the control wells) under microscope (4X objective) for live-dead counting. Appropriate vehicle control containing 0.5%v/v DMSO was also included in the experiment.

### 2.4. Motility assay

Synchronized worms were incubated at 22±1°C with or without extract (600 ppm) in 24-well plates, wherein each well contained approximately 100 worms in 1 mL of M9 buffer. While microscopic observation qualitatively confirmed more agile movement in worm population incubated with extract, quantification of the motility in control vs. experimental wells was achieved employing an automated worm tracker machine, WMicrotracker ARENA (Phylumtech, Argentina). This machine detects the worm locomotion through infrared cameras, wherein magnitude of the scattering of the infrared light by the moving worms crossing the path of light is proportional to the degree of locomotion in worm population. A software algorithm then calculates the number of activity events per time block and its location on the plate. Worm activity counts were recorded once everyday, till progenies appeared in the extract-containing wells (because presence of progenies will contribute to higher activity counts).

### 2.5. Metabolic activity assay

Alamar Blue^®^ assay is widely used to quantify viability or metabolic activity of cells or organisms (Hamid et al., 2004; Tritten et al., 2012), wherein the metabolically active cells reduce the blue dye (resazurin) to a pink or red coloured product (resorufin), and intensity of this colour is proportional to the viability of the biological entity. Synchronized worms were incubated at 22±1°C with or without extract (600 ppm) in 24-well plates, wherein each well contained approximately 100 worms in M9 buffer. One hundred µL of Alamar Blue^®^ (Thermofisher) was added into each well, making the total volume 1 mL, before incubation started. To quantify the amount of dye reduced, after every 24 h, content from wells was transferred into a separate plastic vial (1.5 mL), followed by centrifugation (13,600 g at 25°C) for 10 min. Then the supernatant was read at 570 nm (Agilent Cary 60 UV-vis). This wavelength reads the intensity of the pink colour of the reduced product. Total three wells were set for control as well as experimental worm population, and content from one well, from control and experimental group each, was used on a particular day. Appropriate abiotic controls (containing the dye and other media components, but no worms) were also included in the assay.

### 2.6. Whole Transcriptome Analysis

To unravel the molecular mechanisms through which *W. somnifera* conferred beneficial effect on the worms with respect to lifespan and fertility, the gene expression pattern of extract-treated worms was compared with that of the control worm population at the whole transcriptome scale. While harvesting the worms for RNA isolation on seventh day of experiment (when ∼50% worms in control population were dead), progenies were separated from the experimental population by resting the tubes containing worms incubated with extract in static condition for few minutes, allowing settling down of adult worms. Progenies were removed by collecting the upper layer of liquid. This removed liquid was microscopically observed to confirm presence of largely progeny worms in it, and not the adult ones added at the starting day of the experiment. Following this, worms were washed thrice with sterile M9 buffer before proceeding further. The overall workflow of this whole transcriptome analysis (WTA) aimed at capturing a holistic picture regarding modes of action of the test extract is given in Figure S2.

#### 2.6.1. RNA Extraction

Worm RNA was extracted by the Trizol (Invitrogen Bioservices, Mumbai, India; 343909) method. RNA was precipitated using isopropanol, followed by washing with ethanol (75%), and then the RNA was suspended in nuclease-free water. RNA thus extracted was quantified on Qubit 4.0 fluorimeter (Thermofisher, Mumbai, India; Q33238) making use of an RNA HS assay kit (Thermofisher; Q32851) as per the manufacturer’s protocol. Concentration and purity of RNA were assessed on Nanodrop 1000. RIN (RNA Integrity Number) value was known by assessing the RNA on the TapeStation using HS RNA ScreenTape (Agilent) (Table S1).

#### 2.6.2. Library Preparation

The final libraries prepared using Trueseq standard total RNA (Illumina #15032611) were quantified on a Qubit 4.0 fluorimeter, using a DNA HS assay kit (Thermofisher; Q32851). To determine the insert size of the library, it was queried on Tapestation 4150 (Agilent) using highly sensitive D1000 screentapes (Agilent; 5067-5582). Acquired sizes of both libraries are listed in Table-S1.

#### 2.6.3. Genome Annotation and Functional Analysis

Quality assessment of the raw fastq reads of the sample was achieved using FastQC v.0.11.9 (default parameters) (Andrews 2010). The raw fastq reads were pre-processed using Fastp v.0.20.1 (Chen et al., 2018), followed by a reassessment of the quality using FastQC (Ewels et al., 2016). The processed reads were aligned to the STAR indexed *Caenorhabditis elegans* (Ensembl, WBcel235) genome using STAR aligner v 2.7.9a (Dobin et al., 2013) (parameters: --outSAMtype BAM SortedByCoordinate --outSAMunmapped Within --quantMode TranscriptomeSAM -- outFilterScoreMinOverLread 0.5 --outFilterMatchNminOverLread 0.5 --outSAMattributes Standard). The rRNA and tRNA features were removed from the GTF file of *C. elegans.* The alignment file (sorted BAM) from individual samples was quantified using featureCounts v. 0.46.1 (Liao et al., 2014) based on the filtered GTF file to obtain gene counts. These gene counts for *C. elegans* were used as inputs to edgeR with exactTest (Robinson et al., 2010) for differential expression estimation (parameters: dispersion = 0.1). Gene Ontology (GO) and Kyoto Encyclopedia of Genes and Genomes (KEGG) pathway annotation of the samples was performed using bioDBnet (Mudunuri et al., 2009). The EdgeR exact Test annotated file was filtered based on adjusted *p*-value (FDR) ≤0.05 and Log Fold Change ± 2. Volcano plots were generated using Enhanced Volcano (Blighe et al., 2019), and the MA plots were plotted using ggmaplot function of ggpubr R package (Kassambara et al., 2020). Gene set GO enrichment analysis was done using ShinyGO (Ge et al., 2020) (parameters: Organism: *C. elegans*, FDR≤0.05).

All the raw sequence data were submitted to the Sequence Read Archive. The relevant accession number for control and experimental worm population are SRX20790765 (https://www.ncbi.nlm.nih.gov/sra/SRX20790765) and SRX20790719 (https://www.ncbi.nlm.nih.gov/sra/SRX20790719) respectively.

#### 2.6.4. Network Analysis

From among all the differentially expressed genes (DEG) in *W. somnifera*-exposed *C. elegans*, those fulfilling the dual filter criteria of False Discovery Rate (FDR) ≤ 0.01 and log fold change ≥ 2 were selected for further network analysis. A list of such DEG was input into the STRING (v.12) database (Szklarczyk et al., 2019) to create the PPI (Protein–Protein Interaction) network.

Members of this PPI network were then arranged in descending order of ‘node degree’ (a quantitative indication of connectivity with other proteins or genes), and those above a specified threshold value were forwarded for ranking by the cytoHubba plugin (v.3.9.1) (Chin et al., 2014) of Cytoscape (Shannon et al., 2003). As cytoHubba uses twelve different ranking methods, we considered the DEG top-ranked by a minimum of 6 different methods (50% of the total ranking methods) for further investigation. A separate PPI network of these top-ranked proteins was then generated. The above-described sequence of operations enabled us to end up with a limited number of proteins fulfilling multiple biological and statistical significance criteria simultaneously: (i) FDR ≤ 0.01; (ii) log fold change ≥ 2; (iii) relatively higher node degree; (iv) top-ranking by a minimum of 6 cytoHubba methods.

### 2.7. Polymerase Chain Reaction (RT-PCR)

Differential expression of the potential hubs found using network analysis of DEG revealed from WTA was further confirmed using PCR. Primer designing for the shortlisted genes was carried out using Primer3 Plus (Untergasser et al., 2007). These primer sequences (Table 1) were checked for their binding exclusivity to the target gene sequence within the whole *C. elegans* genome. RNA extraction and quality checks were carried out in the same way as described in the preceding section. cDNA synthesis was carried out using the synthesis kit SuperScript™ VILO™ (Invitrogen Biosciences). PCR assay employed the gene-specific primers purchased from Sigma-Aldrich. The gene bearing ID ‘WBGene00014018’ coding for a RNA binding protein was run as the endogenous control. FastStart Essential DNA Green Master mix (Roche, Darmstadt, Germany; 06402712001) was used as the reaction mix. The real-time PCR assay was performed on a Quant Studio 5 real-time PCR machine (Thermo Fisher Scientific, Waltham, MA, USA). The temperature profile followed is given in Table S2.

**Table 1.**
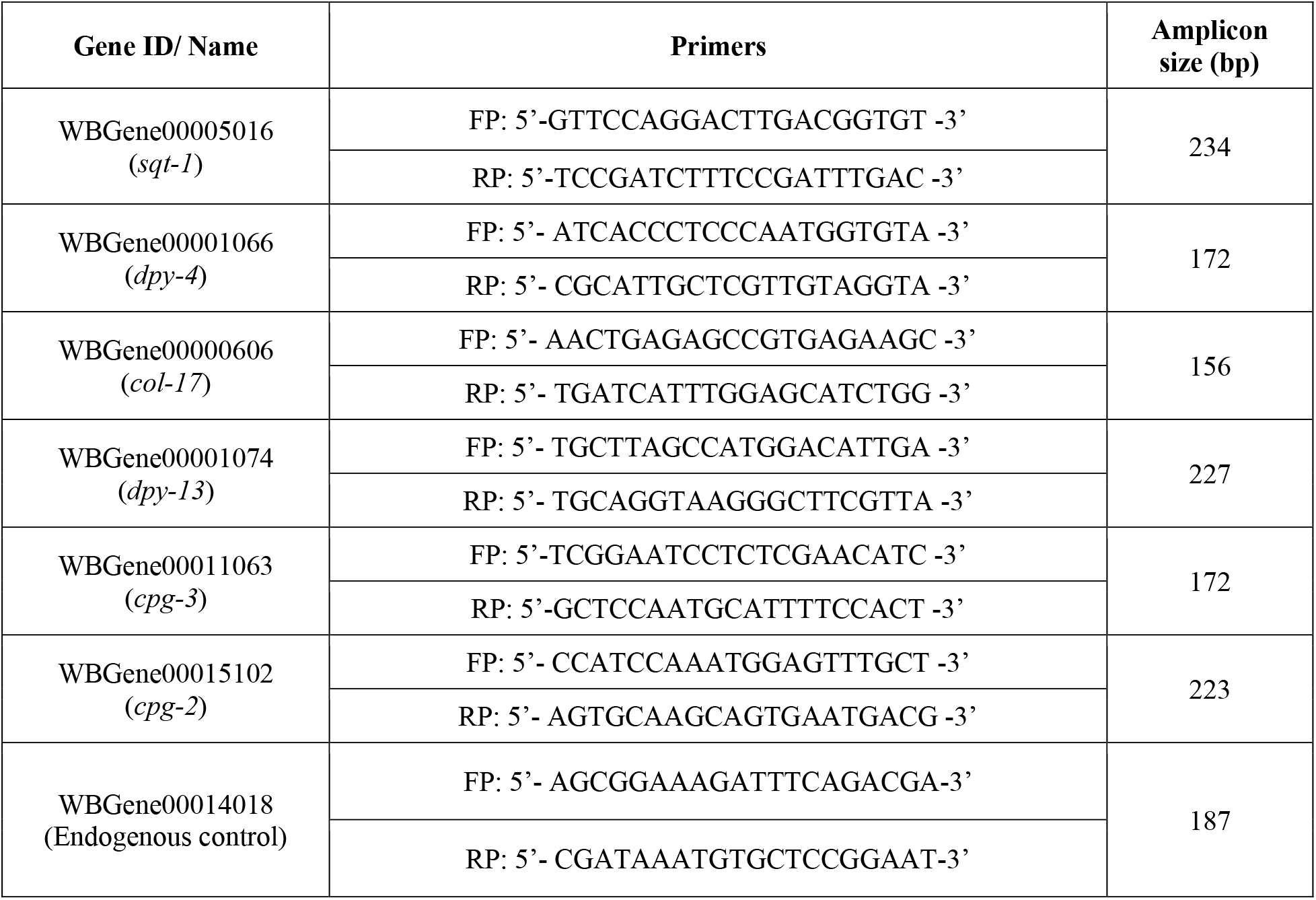
Primer sequences for RT-PCR validation of the selected genes.

**Table 2.** List of DEG satisfying the dual criteria of log fold-change ≥2 and FDR≤0.01.

| Sr. No | Gene ID | Gene | Product/function | Up-/Down-regulation | Log fold change |
| --- | --- | --- | --- | --- | --- |
| 1 | WBGene00006929 | *vit-5* | Vitellogenin-5 | ↑ | 10.78 |
| 2 | WBGene00012592 | Y38E10A.14 | Uncharacterized protein | ↓ | 9.02 |
| 3 | WBGene00011501 | *rmd-1* | Regulator of microtubule dynamics protein 1 | ↑ | 8.06 |
| 4 | WBGene00013391 | Y62H9A.3 | Uncharacterized protein | ↑ | 7.92 |
| 5 | WBGene00019146 | H02F09.3 | Uncharacterized protein | ↓ | 7.75 |
| 6 | WBGene00017501 | *pud-3* | PUD domain-containing protein | ↑ | 5.97 |
| 7 | WBGene00012261 | *lpr-3* | Lipocalin-Related protein | ↓ | 5.74 |
| 8 | WBGene00021005 | *ule-1* | Chitin-binding type-2 domain-containing protein | ↑ | 5.63 |
| 9 | WBGene00012186 | *mlt-11* | Uncharacterized protein | ↓ | 5.52 |
| 10 | WBGene00006926 | *vit-2* | Vitellogenin-2 | ↑ | 5.36 |
| 11 | WBGene00009429 | *irg-5* | CUB like domain-containing protein | ↓ | 5.35 |
| 12 | WBGene00000683 | *col-109* | Cuticle collagen N-terminal domain-containing protein | ↓ | 5.04 |
| 13 | WBGene00019148 | H03E18.1 | Uncharacterized protein | ↓ | 5.04 |
| 14 | WBGene00017498 | *pud-4* | PUD domain-containing protein | ↑ | 4.97 |
| 15 | WBGene00000742 | *col-169* | Cuticle collagen N-terminal domain-containing protein | ↓ | 4.9 |
| 16 | WBGene00022033 | Y65B4BL.1 | Uncharacterized protein | ↓ | 4.9 |
| 17 | WBGene00006930 | *vit-6* | Vitellogenin-6 | ↑ | 4.49 |
| 18 | WBGene00000618 | *col-41* | Cuticle collagen N-terminal domain-containing protein | ↓ | 4.43 |
| 19 | WBGene00018268 | F41C3.2 | Uncharacterized protein | ↓ | 4.4 |
| 20 | WBGene00016636 | *perm-2* | Permeable eggshell | ↑ | 4.35 |
| 21 | WBGene00001074 | *dpy-13* | Cuticle collagen dpy-13 | ↓ | 4.27 |
| 22 | WBGene00009035 | *gfat-2* | Glutamine--fructose-6-phosphate transaminase (isomerizing) | ↑ | 4.19 |
| 23 | WBGene00017892 | F28B4.3 | Uncharacterized protein | ↓ | 4.16 |
| 24 | WBGene00000606 | *col-17* | Collagen | ↓ | 4.04 |
| 25 | WBGene00012468 | Y17G7B.17 | Uncharacterized protein | ↓ | 3.98 |
| 26 | WBGene00022816 | *fbn-1* | Fibrillin homolog | ↓ | 3.81 |
| 27 | WBGene00022103 | *cdh-12* | Cadherin family | ↓ | 3.81 |
| 28 | WBGene00001066 | *dpy-4* | Cuticle collagen N-terminal domain-containing protein | ↓ | 3.78 |
| 29 | WBGene00015102 | *cpg-2* | Chondroitin proteoglycan-2 | ↑ | 3.76 |
| 30 | WBGene00010790 | *sodh-1* | Alcohol dehydrogenase 1 | ↑ | 3.76 |
| 31 | WBGene00006787 | *unc-52* | Basement membrane proteoglycan; Ig-like domain-containing protein | ↓ | 3.7 |
| 32 | WBGene00005018 | *sqt-3* | Cuticle collagen 1 | ↓ | 3.58 |
| 33 | WBGene00011063 | *cpg-3* | Chondroitin proteoglycan 3 | ↑ | 3.54 |
| 34 | WBGene00007709 | *clec-87* | C-type lectin domain-containing protein 87 | ↑ | 3.52 |
| 35 | WBGene00011321 | *fil-1* | Fasting Induced Lipase | ↑ | 3.48 |
| 36 | WBGene00007723 | C25F9.2 | DNA-directed DNA polymerase | ↓ | 3.37 |
| 37 | WBGene00005016 | *sqt-1* | Cuticle collagen sqt-1 | ↓ | 3.37 |
| 38 | WBGene00010070 | *nep-17* | Neprilysin metallopeptidase family | ↓ | 3.28 |
| 39 | WBGene00006436 | *ttn-1* | Titin homolog | ↓ | 3.09 |
| 40 | WBGene00006801 | *unc-68* | Uncharacterized protein | ↓ | 3.08 |
| 41 | WBGene00002915 | *let-805* | Uncharacterized protein | ↓ | 3.07 |
| 42 | WBGene00004130 | *ketn-1* | Uncharacterized protein | ↓ | 3.05 |
| 43 | WBGene00000998 | *dig-1* | Mesocentin | ↓ | 3 |
| 44 | WBGene00006876 | *vab-10* | GAR domain-containing protein | ↓ | 2.89 |
| 45 | WBGene00220238 | ZK185.9 | Uncharacterized protein | ↑ | 2.74 |

Genes in this table are arranged in decreasing order of fold-change value.

### 2.8. Statistics

All values reported are derived from three or more independent experiments, wherein each experiment contained three replicates (unless specified otherwise). Statistical significance was assessed using a t-test performed in Microsoft Excel^®^ (Version 2016), and data with p ≤ 0.05 were considered to be significant

## 3. Results and Discussion

### 3.1. Worms incubated with the root extract exhibited extended lifespan as well as higher fertility than control population

We incubated worms with the water-soluble fraction of the root extract or that dissolved in DMSO, and observed the worms over a period of 12 days (till almost all worms in control wells died) for live-dead count, morphology, agility, and whether any progenies are formed. While comparison between control and experimental wells was made on daily basis, two time-points can particularly be considered important i.e. the days on which control population exhibited ∼50% and ∼100% death. With respect to these two endpoints, while the DMSO-dissolved extract could impart a longevity benefit to the worms at all tested concentrations ≥100 µg/mL, water-soluble fraction of the extract could do so at ≥250 µg/mL. Additionally, concentrations ≥ 500 µg/mL of both DMSO-solubilized and water-solubilized extract supported worm fertility from day-4 onward, as evident from appearance of progenies in the extract-containing wells, and their absence in control wells (Videos S1-S2). Till day-12 these progenies could be differentiated from the parent worms based on size. Thereafter almost all the parent worms in control wells were dead, and size of the surviving parent worms and their progenies in the experimental wells became similar to the extent that they could not be differentiated. The magnitude of survival benefit (i.e. higher number of surviving worms in extract-containing wells than control wells) conferred on the worms by DMSO-solubilized extract (Figure 1A) was somewhat higher than that of water-solubilized extract (Figure 1B), except at 750 µg/mL. At the latter concentration, the water-solubilized fraction of the extract performed better (p=0.0004) than the DMSO-solubilized extract, as per the final-day endpoint. We decided to perform further experiments with water-solubilized fraction only, as water-solubility of any therapeutic preparation is looked favourably with respect to bioavailability (Bhalani et al., 2022).

**Figure 1.**
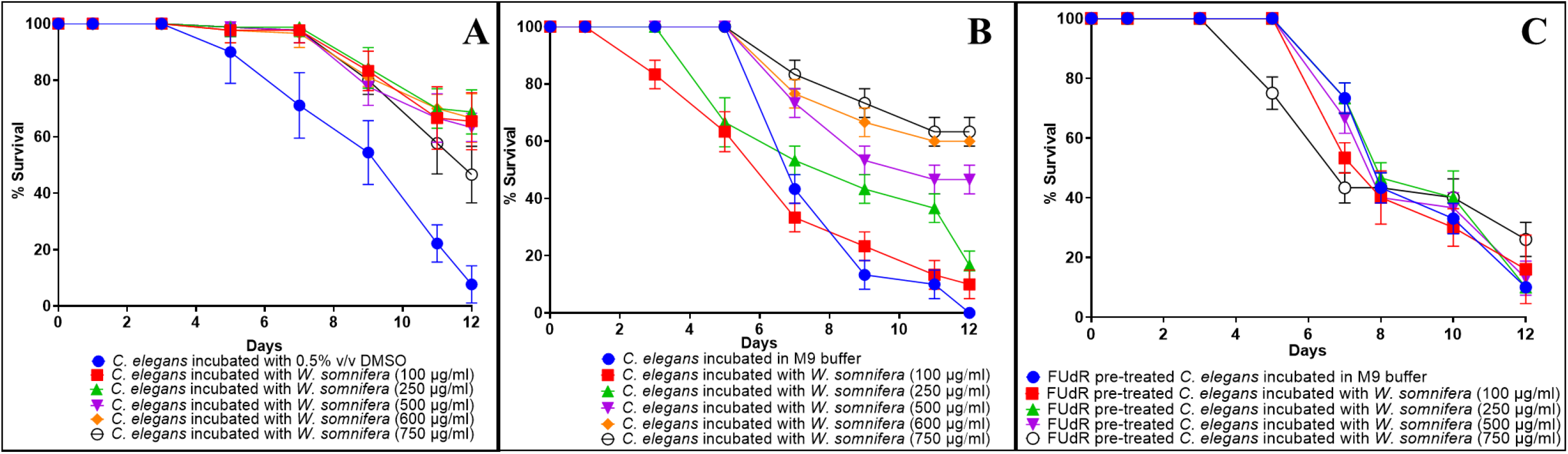
*Withania somnifera* imparts longevity extension on *C. elegans*. All worms in control wells (i.e. worms not fed with extract) were dead by the 12^th^ day. Concentrations ≥500 µg/mL of both DMSO-solubilized and water-solubilized extract supported worm fertility from day-4 onward. Bigger body size, higher motility, and progeny formation in presence of the extract in experimental wells can be visualized in supplementary videos S1-S4. DMSO (0.5%v/v) or FuDR (15mM/mL of NGM agar) did not affect worm lifespan. Only selected concentrations are displayed in this figure to avoid overcrowding of graphs. (A) Worms fed with DMSO-solubilized *W. somnifera* extract at 100, 250, 500, 600, 750 μg/mL scored 65.55%*** ± 10.13, 68.88%*** ± 7.81, 63.33%*** ± 5, 66.66%*** ± 8.66 and 46.66%*** ± 10 better survival on the 12^th^ day compared to control. (B) Worms fed with water-solubilized *W. somnifera* at 100, 250, 500, 600, 750 μg/mL scored 10%*** ± 5, 16.66%*** ± 5, 46.66%*** ± 5, 60%*** ± 0 and 63.33%*** ± 5 better survival on the 12^th^ day compared to control. (C) FuDR pre-treated worms fed with water-solubilized *W. somnifera* extract did not display any extended lifespan except at 750 μg/mL (26.66%*** ± 5.16 better survival on the 12^th^ day compared to control. *** p ≤ 0.001

While the results described in Figure1A-1B demonstrated the beneficial effect of test extract on worm lifespan as well as fertility, to investigate its effect on worm longevity separately from fertility, we repeated this assay using FUdR pre-treated worms, wherein eggs were allowed to hatch on FUdR-containing plates pre-seeded with *E. coli* OP50, and then resulting L3-L4 stage worms were washed with M9 buffer before being transferred to a fresh NGM agar plate for further use. FUdR (5-fluoro-20 -deoxyuridine; HiMedia) can sterilize the adult nematode worms without affecting their development (Mitchell et al., 1979). However, FUdR-pre-treated worms could not benefit from the pro-longevity effect of water-soluble fraction of the *W. somnifera* root extract except at 750 µg/mL on day-12 (Figure 1C), and here too the effect was much lesser than when FUdR was not used (Figure 1B). These results raise a caution regarding use of FUdR in lifespan assays with *C. elegans*, as this may lead to false-negative conclusions. Though the practice of using FUdR-treated worms is popular among the worm researchers studying longevity and aging, in order to avoid the labour intensive separation of offsprings from adults during the reproductive period, our results show that FUdR can prevent detection of pro-longevity activity in a bioactive extract, even when the said activity is there. This corroborates with the earlier suggestion by Aitlhadj and Stürzenbaum (2010), that the effect of FUdR is not neutral and owing to its mechanism of actions (inhibition of DNA and RNA synthesis, death of mitotic cells, and the inhibition of protein synthesis), its inclusion in the assays may result in misinterpretation. Hence it can be suggested that while investigating any natural product’s biological effect in *C. elegans* model, it is more useful to do wholistic assay assessing multiple parameters like lifespan, health, metabolic activity, fertility, etc., rather than doing an assay exclusively focusing on longevity using additional chemicals like FUdR which may prove to be a confounding factor.

### 3.2. *W. somnifera* root extract positively affects worm motility and metabolic activity

Worms incubated in presence of the test extract exhibited more active movement (Figure 2A), which can be considered another indicator of overall good health. This observation matches well with the high fertility of extract-exposed worms described in preceding section, since the egg-laying active state and the defecation motor program are both linked to changes in forward and reverse locomotion (Hardaker et al., 2001; Nagy et al., 2015). Automated monitoring of *C. elegans* movement is a useful and faster healthspan-based method to study aging (Zavango et al., 2024). Besides positively impacting worm lifespan, fertility, and motility, the root extract also had a stimulatory effect on work metabolic activity (Figure 2B). Worms incubated in presence of the extract were found to have higher reducing potential (captured in terms of their ability to reduce the dye Alamar blue), which can be taken as an indication of better health, as healthy living cells maintain a reducing state within their cytosol (Longhin et al., 2022).

**Figure 2.**
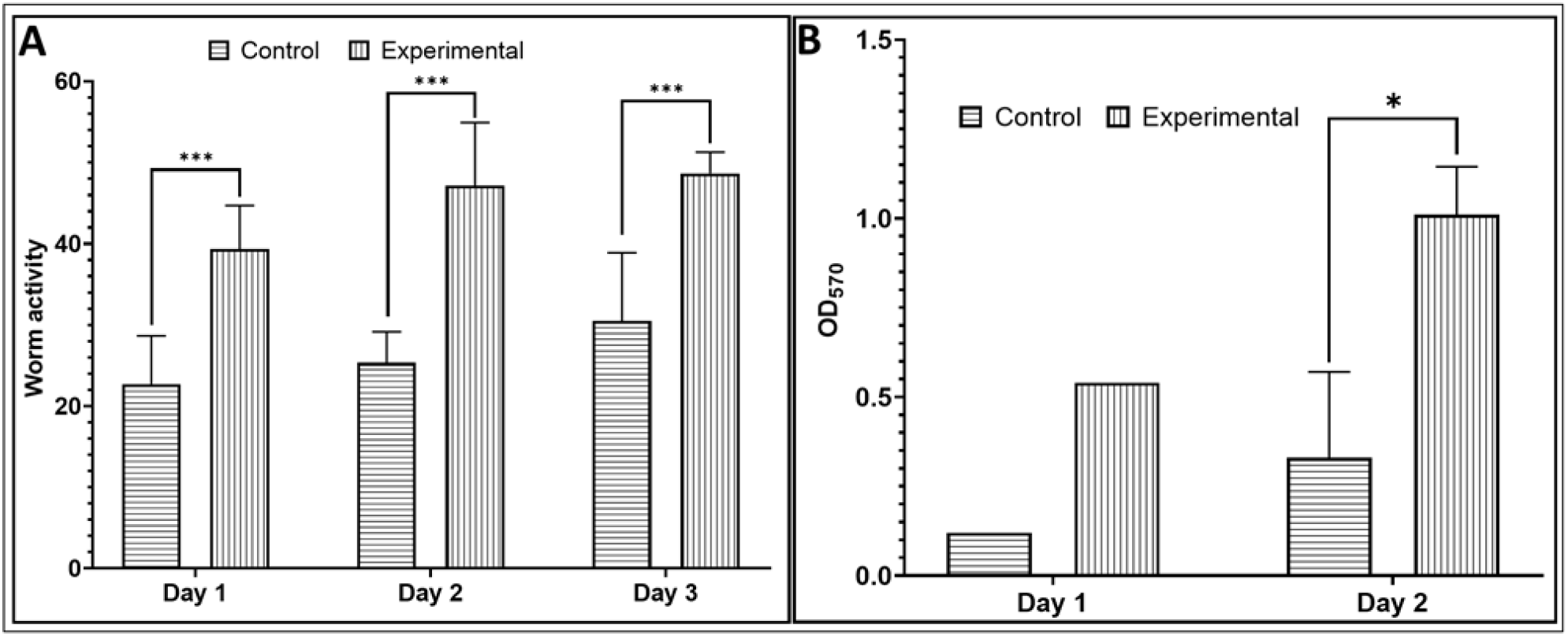
*W. somnifera* root extract has a positive effect on worm motility. **(A)** and metabolic activity **(B)**. Worm activity plotted on y-axis in figure (A) is quantification of their movement as measured by an automated worm tracker. Higher motility in extract-treated worms plotted here is in line with the visual observation under microscope. OD570 in figure (B) corresponds to the amount of dye reduced by metabolic activity of the worm population. * p ≤ 0.05, *** p ≤ 0.001

Overall, the root extract imparted its beneficial effect on multiple health parameters of the worm i.e. lifespan, fertility, motility, and metabolic activity. Increased egg laying in extract-exposed older adults reflects an increase in the number of eggs expelled per vulval opening, as well as, longer active behavior states. Since the timing of expulsive behaviors (defecation and egg laying) is regulated by sensory mechanisms that detect changes in internal pressure and/or stretch to maintain homeostasis (Ravi et al., 2019), we believe that the test extract had a multifactorial impact on worm physiology at different levels. Feedback of successful egg laying in presence of the extract might have signaled to the germ line to continue production of oocytes for fertilization. A potential trade-off between reproductive capacity and somatic maintenance in *C. elegans* has already been mentioned in earlier published literature. Higher levels of progeny production is a biomarker for studying aging and correlates positively with a longer lifespan (Klass, 1977; Pickett et al., 2013). Prolonged reproduction can have a beneficial impact on lifespan.

Since most promising effect of the root extract on worm’s overall health (i.e. lifespan and fertility) were observed while using water-soluble fraction of the extract, and the maximum beneficial effect was observed at 600-750 µg/mL, we decided to investigate the gene expression pattern of worms exposed to the test extract at 600 µg/mL at the whole transcriptome level. Since the biological effect of both these concentrations (600 and 750 µg/mL) was statistically similar (p>0.05), we went ahead with 600 µg/mL. Before heading for transcriptome analysis, additional confirmatory experiments were also done to have more confidence in beneficiary potential of *W. somnifera,* whose results are described below.

### 3.3. *W. somnifera* root extract exerts its beneficial effect on the nematode worm by triggering differential expression of multiple genes

A whole transcriptome level comparison of gene expression profile of the extract-treated worms with their extract-non-exposed counterparts revealed all the DEG in the experimental worm population. Keeping the criteria of log FC ≥1.5 and FDR (false discovery rate) ≤0.05 the differentially expressed gene (DEG) count was 85. However, to have higher confidence in our data interpretation, we set a more stringent dual criteria of log FC ≥2 and FDR≤0.01 to shortlist the genes for further analysis. A total of 16 upregulated and 29 downregulated DEG passing the said dual criteria are listed in Table-2. These 45 DEG comprises 0.83% of total *C. elegans* genome (∼5423 genes). Function-wise categorization of these DEG is presented in Figure-3, and corresponding heat map (Figure S3) and volcano plot (Figure S4) is provided in supplementary file.

Log fold-change values of the DEG ranged from 2.74-10.78. Among the top-17 DEG, three of the upregulated genes coded for different vitellogenin lipoproteins. In *C. elegans*, the synthesis of the yolk lipoprotein (vitellogenin) occurs in the endoplasmic reticulum of the intestine, and is initiated before the worm achieves adulthood. Since vitellogenesis persists throughout and beyond the reproductive stage, lipoproteins accumulate in the passive circulatory system found between tissues (pseudocoelom) as the animal ages. While Seah et al. (2016) reported overexpression of the vitellogenin to be associated with a reduction in the lifespan of worms, our study has found an upregulation of *vit* genes in extract-exposed worms experiencing a longer life. Increased fertility observed in our study in *W. somnifera*-exposed worms corroborates well with the primary function of abundant vitellogenesis in *C. elegans* proposed by Perez and Lehner (2019), that is to support post-embryonic development and fertility, especially in harsh conditions (in our study, the ageing gnotobiotic worms faced starvation, as they were not provided any bacterial food during the whole assay). Vitellogenins are a family of yolk proteins that are the most abundant among oviparous animals. In *C. elegans*, the six vitellogenins are among the most highly expressed genes in the adult hermaphrodite intestine, which contributes towards ensuring yolk for eggs. Vitellogenins can act as an intergenerational signal mediating the influence of parental physiology on progeny. While *vit* genes are known to be important in regulating lifespan of the worms (Sornda et al., 2019), our results indicate that vitellogenins may play other important roles as well in worm physiology. Since vitellogenins are widely present across the tree of life, *W. somnifera’*s pro-health effect mediated in part through upregulation of *vit* genes can be expected to be relevant for other forms of life including humans. On the basis of sequence similarity, vitellogenins are suggested to be the ancestor of human apoB, the principal component of low density lipoprotein (Baker, 1988). Interestingly the three upregulated *vit* genes in our transcriptome data belong to three different groups of the vitellogenin family. Of them, the *vit*-6 is the only member of its group, and this protein was found by Nakamura et al. (1999) as a major carbonylated protein in aged *C. elegans*, with a potential role in protecting other cellular components from oxidative stress. Such oxidative stress can be believed to be experienced by the worms in our study owing to starvation (Tao et al., 2017). Vitellogenin has recently been shown to play non-nutritional roles too, by functioning as an immunocomponent factor and antioxidant (Li and Zhang, 2017).

Upregulation of above mentioned *vit* genes gets justified in view of upregulation of multiple other genes (*cpg-2, cpg-3, ule-1* and *perm-2*) too, contributing to the common function of the hierarchical assembly of the eggshell and permeability barrier in this worm. The *C. elegans* eggshell is composed of an outer vitelline layer, a middle chitin layer, and an inner layer of chondroitin proteoglycans. Since *W. somnifera* extract promoted fertility of worms in our study, the genes contributing towards formation of the eggshell can be expected to be upregulated. Among products coded by such genes, cpg-2 is required for eggshell impermeability, and possibly an important cargo. Upregulation of *ule-1* (chitin binding domain protein) and the chondroitin proteoglycans (*cpg-2* and *cpg-3)* can be understood in light of the fact that chitin and the *cpg-1/2* localize to the middle and inner layers of the trilaminar eggshell, respectively (Olson et al., 2012). Simultaneous upregulation of these genes can be explained by the fact that formation of the inner chondroitin proteoglycan (CPG) layer requires prior deposition of the chitin layer. The chondroitin chains play essential roles in embryonic development and vulval morphogenesis of worms. *cpg-2* is among those proteoglycans on which embryonic cell division is dependent. Owing to its expression during embryonic development and chitin-binding property, it has been suggested to have a structural role in the egg. (Olson et al., 2006). Another member of the cpg group in our list of DEG, cpg-3 is also implicated in chitin and chondroitin biosynthesis and eggshell formation (Kudron et al., 2013). Upregulation of a uterine protein (ule-1) in extract-fed fertile worms can be correlated to the fact that uterine proteins turn over rapidly in young worms than old, and the extract-fed worm population was indeed displaying more ‘youthful’ behaviour in terms of active motility and reproduction on the day of worm collection for transcriptome purpose. Since uterine proteins are removed by egg-laying in young worms, their upregulation may be compensated by the worms through active reproduction. If that is not the case, then upregulation of *ule* genes may shorten the lifespan (Zimmerman et al., 2015). Upregulation of the *ule-1* gene in our extract-fed actively reproducing worm population matches with earlier published role of this gene, wherein it was identified in mass RNAi screens to be required for normal progeny production (Maeda et al., 2001; Green et al., 2011), and its knockdown marginally reduced the brood size. One of the upregulated genes in extract-fed worms, *perm-2* is among the key components of the vitelline layer structural scaffold, whose depletion from the scaffold can lead to a porous vitelline layer which permits soluble factors to leak through the eggshell resulting in embryonic death (González et al., 2018). perm-2 is among those proteins which are required for eggshell integrity, and its upregulation corroborates well with increased fertility of the *W. somnifera*-fed worms.

Among other upregulated genes in extract-treated worms were two *pud* genes, *pud-3* and *pud-4.* The PUD gene family comprises of six genes located next to each other on chromosome V. Pud-genes are unique to *Caenorhabditis* spp. without known orthologues in other nematodes While upregulation of other *pud* genes (*pud-1* and *pud-2*) has been reported earlier (Ding et al., 2013) in long-lived *C. elegans,* our study found different genes of the same group to be upregulated in the experimental worms. Of them, *pud-3* is known to get expressed in the nuclei of the largest hypodermal cell (hyp7) which wraps around most of the worm body, and also in the rectal gland cells, pharyngeal muscle pm3, and less frequently in the intestine and a few neuron-like cells in the head. *pud-4* gets expressed most strongly in the hyp7 nuclei and sporadically in the intestine. The nuclear localization of the PUD proteins (including those upregulated in our study) is suggested as an indication of their involvement in regulation of the transcription of the collagen genes. Slow growth of *C. elegans* after *pud-4* knockdown in worms reported by Cui et al. (2007) corroborates with our study finding the same gene getting upregulated in fast growing *W. somnifera*-exposed worms. *pud-3* and *pud-4* were downregulated in their study in worms exposed to the anthelmintic drug ivermectin. It may be thought that these *pud* genes are upregulated under favourable conditions, and downregulated under difficult conditions.

Of the top-24 DEG, four (*col-109, col-169, col-41*, and *col-17*) belonged to the *col* group of genes, and all these four were downregulated in extract-exposed worms. Downregulation of this *col* group of genes corroborates well with simultaneous downregulation of other genes (*sqt-1, sqt- 3, dpy-4,* and *dpy-13*) contributing to same function e.g. cuticle collagen synthesis. These collagen genes are involved in the formation and maintenance of the worm cuticle. Collagens are essential for providing structural support and elasticity to various tissues, and hence are crucial for the integrity and protective function of the worm’s outer layer. Cuticle collagens, the major cuticle component, are encoded by a large family of *col* genes and, interestingly, many of these genes express predominantly at a single developmental stage (Abete-Luzi and Eisenmann, 2018). The nematode’s genome encodes 177 collagens (Sandhu et al., 2021). It is possible that the worm regulates their expression to modulate cuticle permeability as a part of its adaptive response to external environment. Outer covering of the nematode *C. elegans*, the cuticle, is synthesized five times during the worm’s life by the underlying hypodermis. It is possible that since the life of worms exposed to the root extract in our study was prolonged, they reduced the pace of cuticle synthesis by downregulating expression of certain *col* genes specifically important for that particular stage of life cycle. It is possible that extract-induced longevity and pro-fertility effect might have caused the worms to undergo a rearrangement of their cuticle microstructure and permeability, may be in order to allow enhanced entry of the beneficial phytocompounds. Apical extracellular matrices (aECMs) are complex extracellular compartments that form important interfaces between animals and their environment. In the adult *C. elegans* cuticle, layers are connected by regularly spaced columnar structures (Adams et al., 2023). It is possible that the extract would have triggered remodeling of this precise aECM patterning at the nanoscale. While multiple *col* genes were downregulated in extract-treated worms, since not much is known about role of these genes in maintaining the structure or barrier function of the cuticle and the cuticle permeability, it is difficult to discuss how precisely downregulation of these genes can be correlated to enhanced lifespan and fertility in extract-fed worms.

In total, 8 genes contributing towards collagen synthesis were downregulated in our experimental worm population. This might have led to some degree of collagen restructuring, which in turn may have dramatic effects on organism’s morphology. Among collagen synthesis-associated genes other than the *col* genes found among our DEG, the *sqt-1* plays a unique role in determining the structure of the cuticle and the morphology of the animal. This gene has been reported to encode a collagen, which is critical for morphogenesis of *C. elegans* (Kramer et al., 1988). Other genes belonging to same group, *sqt-3* codes for an early cuticle collagen. Its downregulation may have slowed down the formation of the basal striated layer of the L1 cuticle, which forms only after embryo elongation (Birnbaum et al., 2023). The C terminus of collagen sqt-3 has been reported for its complex and essential role in nematode collagen assembly (Novelli et al., 2006). Though *sqt-3* is recognized mainly as a component of the cuticle, but other important roles of it have also been indicated. For example, it has been shown to direct anterior–posterior migration of the Q Neuroblasts in *C. elegans* (Lang and Lundquist, 2021). Two more genes coding for cuticle collagen components, *dpy-13* and *dpy-4* were also downregulated in extract-treated worms. Of them, *dpy-13* is among those having specific roles in the formation of the exoskeleton (Nyström et al., 2002), and it has also been identified as a collagen gene influencing the nematode body shape (von Mende et al., 1988). Extract-treated worms in our study experiencing downregulation of *dpy-4* can still be believed to have no cuticle abnormality, as *dpy-4* non-expressing mutant of *C. elegans* were shown to possess normal ultrastructure of the cuticle similar to wild type control animals. The *dpy-4* RNAi animals were shown to bear a periodic arrangement of annuli and furrows, without any irregular indentations (Sandhu et al., 2021).

Two *unc* genes, *unc-52* and *unc-68* were downregulated in extract-exposed worm population. The *unc-52* gene of *C. elegans* codes for a homologue of the basement membrane heparan sulfate proteoglycan perlecan, which was shown to affect gonadal leader cell migrations in hermaphrodite worms through alterations in growth factor signaling. *unc-52* has been proposed to play dual roles in *C. elegans* larval development in the maintenance of muscle structure, as well as, the regulation of growth factor-like signaling pathways (Merz et al., 2003). Downregulation of *unc-52* can be expected to have altered the formation or maintenance of the muscle myofilament lattice (Rogalski et al., 2001). Another downregulated gene, *unc-68* codes for a ryanodine receptor involved in regulating worm body-wall muscle contraction. Ryanodine receptors are not believed to be essential for excitation-contraction coupling in nematodes, but they do act to amplify a calcium signal to the extent sufficient for contraction. Though *unc-68* mutants were reported to be somewhat defective with respect to motility and pharyngeal pumping (Maryon et al., 1996), its downregulation in our extract-exposed worms did not have any negative effect on motility. In fact, they were more actively motile than control worms (Figure 2A; Videos S1-S2). Any possible effect of *unc-68* downregulation on pharyngeal pumping is not of much relevance to our study, as we did not provide any solid (e.g. bacterial cells) food to the worms during the assay. Higher proficiency of *unc-68* mutants with respect to egg laying reported by Maryon et al. corroborates with our observation of increased fertility in extract-exposed worms.

Fold-change wise, the third most differently expressed gene in extract-fed worms was *rmd- 1*, which codes for a microtubule-associated protein, having function in chromosome segregation in *C. elegans*. Importance of the upregulation of this regulator of microtubule dynamics in *W. somnifera*-exposed worms can be understood from the fact that depletion of this protein can induce severe defects in chromosome segregation. Relevance of this result can be visualized better in light of the fact that human homologues of *rmd-1* also bind microtubules, suggesting a function for these proteins in chromosome segregation during mitosis in other organisms as well (Oishi et al., 2007). Non-expression of *rmd-1* leads to defects in chromosome segregation and microtubule outgrowth. It can be said that in addition to regulating microtubule outgrowth, *rmd-1* has specific functions in the execution of proper chromosome segregation, may be by preventing abnormal attachments. *rmd-1,2,3,6* is a family of homologous proteins conserved between humans and *C. elegans,* and perhaps *rmd-1* is the most important of them all, as paternal mitochondria from an *rmd-2, rmd-3*, *rmd-6* triple mutant were reported to be properly positioned in the *C. elegans* zygote, with a possible explanation that *rmd-1* carries out all the essential functions of this gene family (Juanico et al., 2021).

The seventh top differently expressed gene in experimental worm population was that coding for a lipocalinprotein *lpr-3*, which was downregulated 5.74-fold. Lipocalins in general are required for apical extracellular matrix organization and remodeling in *C. elegans. lpr-3* has a distinct function in precuticular glycocalyx of developing external epithelia. Experimental worms under the influence of root extract experiencing a downregulation of this gene can be expected to have altered phenotype with respect to maintenance of a passable excretory duct lumen, eggshell degradation, shedding of cuticle during molting, cuticle barrier function, and clearance and remodeling of apical extracellular matrix. Its altered expression would have affected multiple aspects of later cuticle structure or molting in worms. *lpr-3* contributes towards maintenance of a uniform duct lumen diameter, as well as, tube patency during lumen elongation and narrowing. Since LPR-3 is present within a juvenile glycocalyx layer, and the connection between this layer and the mature cuticle is weakened during molting, its downregulation would have affected molting in extract-fed worms. Lipocalins are believed to exert direct effects on aECM lipid, metalloproteinase, and/or glycoprotein content, and thereby affect molting and hatching (Forman-Rubinsky et al., 2017). Downregulation of *lpr-3* in long-lived fertile worms in the extract-fed wells becomes more relevant considering that accumulation of this proteins needs to be prevented for ensuring proper localization of the Zona Pellucida domain protein *let-653* within the vulva precuticle (Cohen et al., 2021). Downregulation of *lpr-3* makes more sense when looked together with downregulation of another molting factor (*mlt-11*) in our long-lived worms. Both these genes are among top-10 differently expressed genes. *mlt-11* is a molting factor, and a putative protease inhibitor in *C. elegans*, necessary for embryogenesis, that localizes to lysosomes and the cuticle. It has been demonstrated to be a secreted protein, which localizes in the cuticle and in a punctate pattern reminiscent of lysosome (Clancy et al., 2023). *mlt-11* is proposed to be acting in the aECM to coordinate remodeling and timely ecdysis. Its downregulation can be expected to influence developmental pace, motility, apolysis, and cuticle structure (Ragle et al., 2022).

Another among top-downregulated genes was *irg-5*, which is an immune effector known to be induced in response to bacterial pathogen attack, as well as by exposure to immunostimulatory compounds (Anderson et al., 2019). Its downregulation may not have any negative effect on the worm lifespan as knockdown of *irg-5* was reported not to shorten the lifespan of *C. elegans* grown on *E. coli* OP50 (Peterson et al., 2019).

Since the transcriptome data has identified multiple genes associated with worm exoskeleton components and muscle structure as differently expressed in extract-fed worms, it is clear that the root extract used in the present study has caused restructuring of the exoskeleton in such a way that age-associated development of stiffness in worm skeleton is delayed, and the worm can age in a healthier fashion. We had conducted transcriptome profiling from the worms after seven days of extract-exposure, but the classic signs of old age and fertility loss usually observed in worms at this stage of life cycle did not yet arise in presence of the extract. Hence *W. somnifera* root extract can be concluded to be capable of delaying ageing and senescence. Under routine conditions, at day seven of adulthood, sarcopenia is apparent in histologically examined worms, which leads to the behavioural change in terms of reduced motor activity. Reduced motility stems from stiffening and thickening of the cuticle, which itself results from unregulated collagen biosynthesis (Herndon et al., 2002), and the root extract used in this study seems to have affected collagen synthesis in such a way that stiffening/ thickening of the cuticle was delayed in the extract-fed worms, and thereby maintaining them in an active motility state for longer period.

### 3.4. Identifying the most important DEG through network analysis

To identify the proteins with high network centrality i.e. hub proteins from among the DEG, we generated protein-protein interaction (PPI) network of upregulated (Figure 4A) and downregulated (Figure 5A) genes separately. The PPI network for the upregulated genes contained 15 nodes connected through 28 edges, with an average node degree of 3.73. Since the number of edges (28) in this PPI network is 28-fold higher than expected (01), this network can be believed to possess significantly more interactions among the member proteins than what can be anticipated for a random set of proteins of same sample size and degree distribution. We arranged members of this PPI network in decreasing order of node degree (Table S3), and those with a score of ≥5 were subjected to ranking by different cytoHubba methods. Then we looked for genes which appeared among the top-5 ranked candidates by ≥6 cytoHubba methods, and 5 such shortlisted genes (Table 3) were further checked for interactions among themselves followed by cluster analysis (Figure 4B), which showed them to be networked with an average node degree score of 2.4. This network possessed more edges (06) as against expected (0) for any similar random set of proteins. The PPI network generated through STRING showed these 5 important genes to be distributed among three different local network clusters. Since *cpg-3* appeared to be part of two different clusters, and both the *cpg* genes were having multiple edges connecting them together (node degree score was 3 for both of them), we selected these two genes (*cpg-2* and *cpg-3*) for further RT-PCR validation. PCR assay did confirm upregulation of these two proteins in extract-exposed worms (Figure 6A).

**Figure 3.**
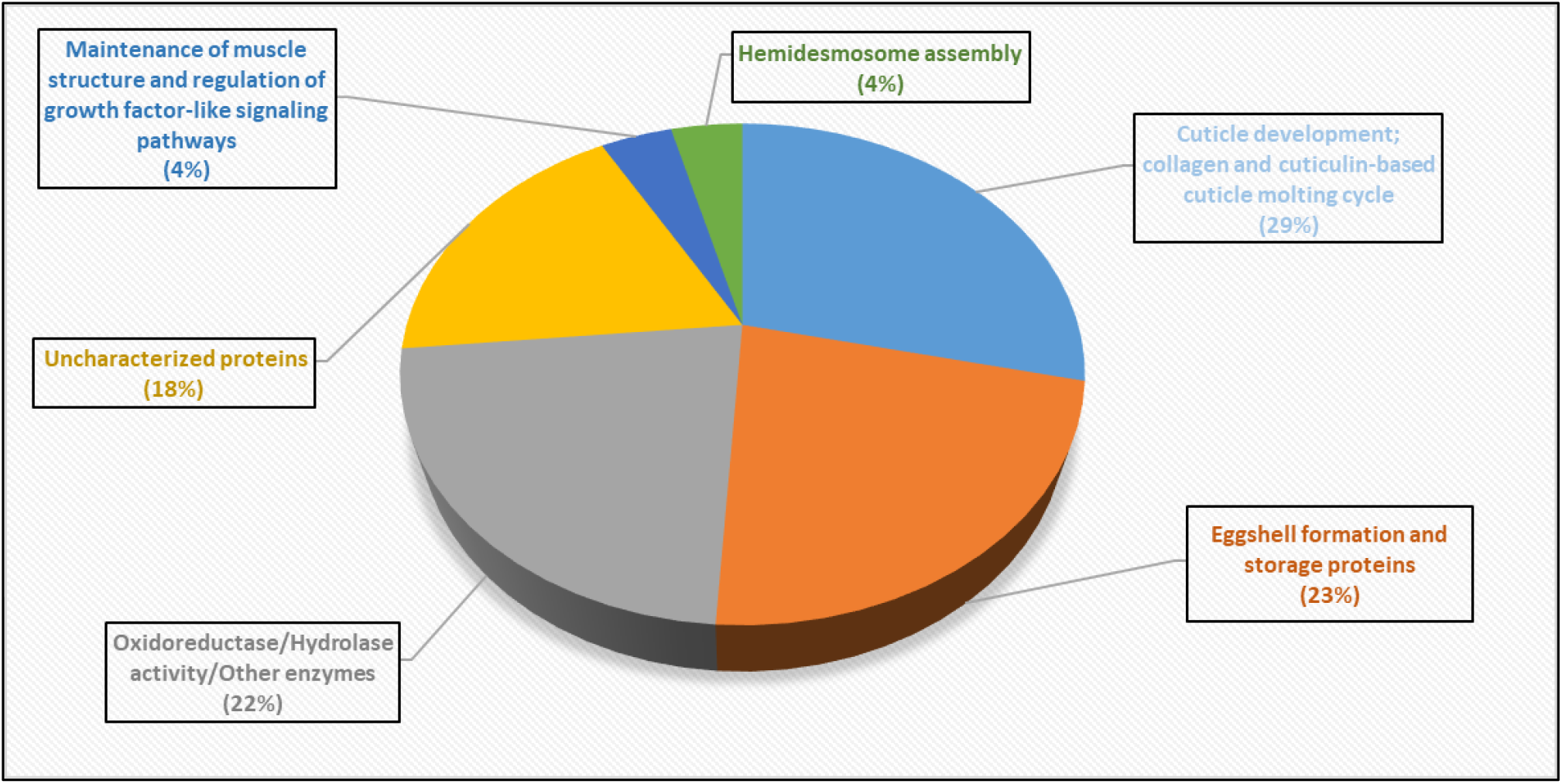
Function-wise categorization of the differentially expressed genes in *W. somnifera-*exposed *C. elegans.* Genes contributing to more than one function are considered in any one category only. Values in parentheses are % of the total DEG.

**Figure 4.**
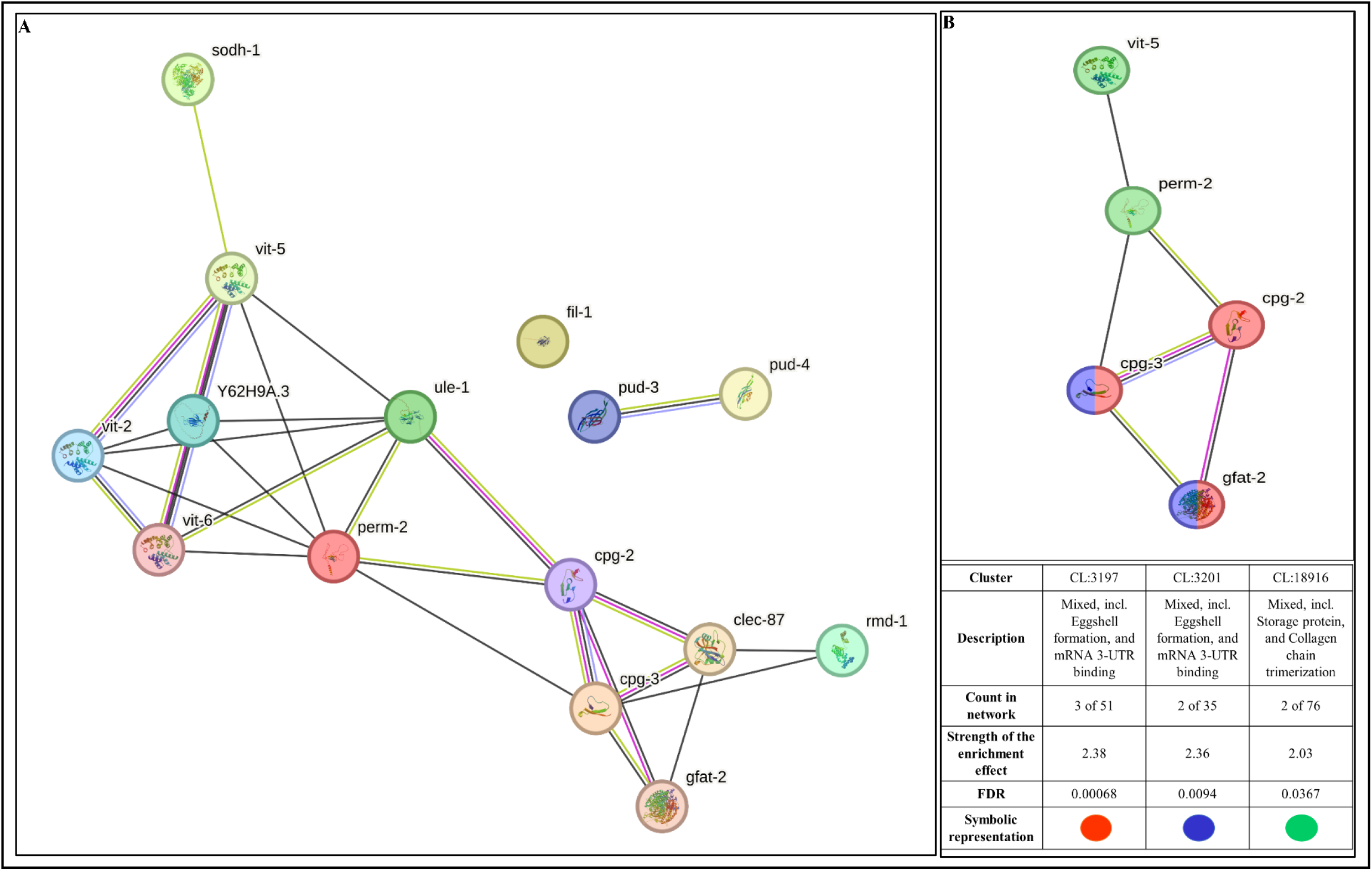
**(A) Protein–Protein Interaction (PPI) network of upregulated genes following the dual criteria of fold change ≥ log 2 and FDR ≤ 0.01 in *W. somnifera*-exposed *C. elegans*.** PPI enrichment p-value 1.0e-16. (https://string-db.org/cgi/network?taskId=badc9unayQGh&sessionId=bmZFhfliKMsM). **(B) PPI network of top-ranked genes shortlisted using cytoHubba among upregulated genes in *W. somnifera-*exposed *C. elegans*.** Value of node degree score for *cpg-2*, *cpg-3*, and *perm-2* is 3, and that for *gfat-2* and *vit-5*, 2 and 1 respectively. PPI enrichment p-value 3.24e-10 (https://string-db.org/cgi/network?taskId=bbPybwRHQ9La&sessionId=bmZFhfliKMsM)

**Figure 5.**
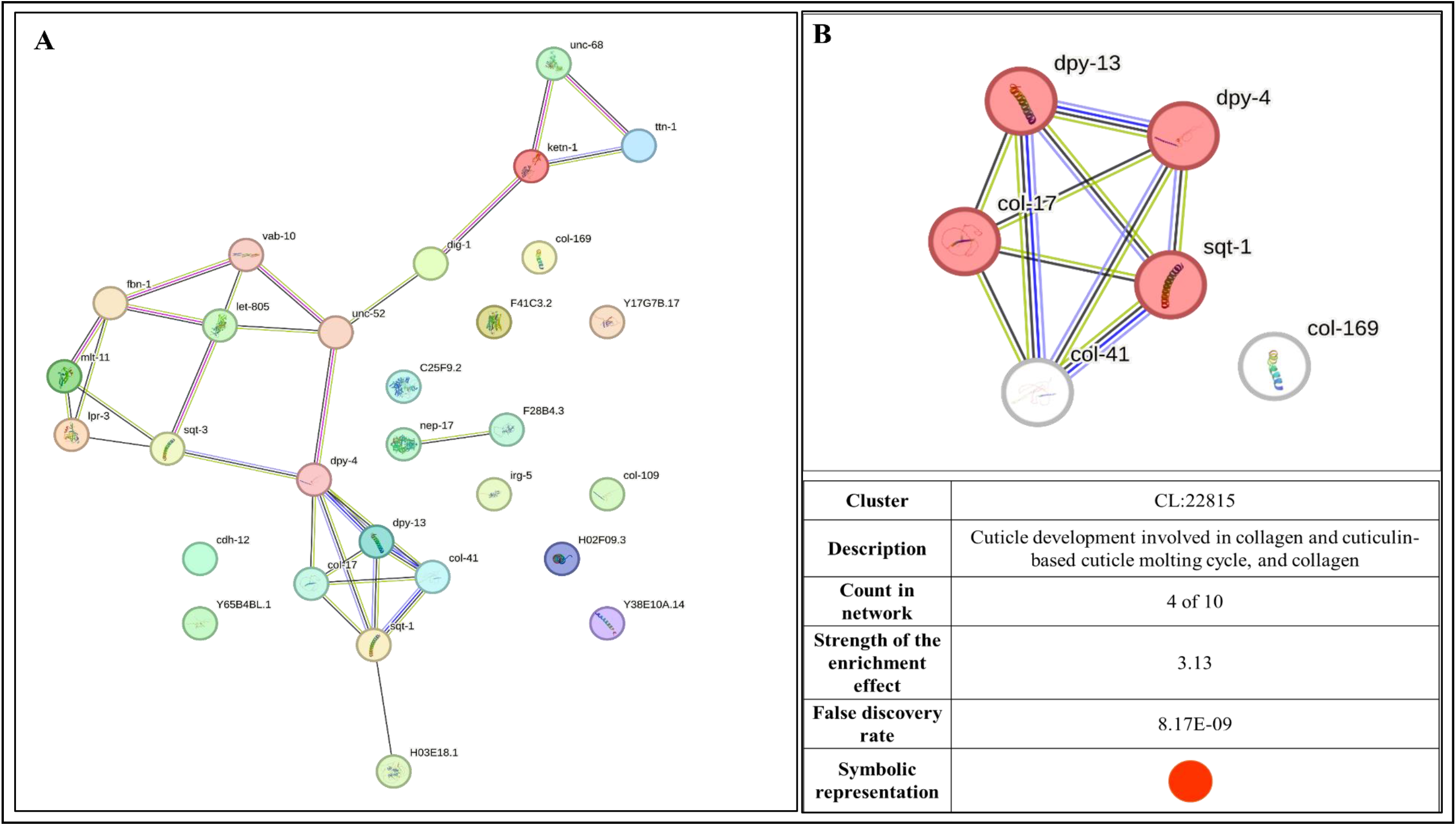
**(A) Protein–Protein Interaction (PPI) network of downregulated genes following the dual criteria of fold change ≥ log 2 and FDR ≤ 0.01 in *W. somnifera*-exposed *C. elegans*.** PPI enrichment p-value 1.0e-16 (https://string-db.org/cgi/network?taskId=bRhNGHhVNwKu&sessionId=bmZFhfliKMsM). **(B) PPI network of top-ranked genes shortlisted using cytoHubba among downregulated genes in *W. somnifera-*exposed *C. elegans*.** PPI enrichment p-value 1.0e-16. (https://string-db.org/cgi/network?taskId=bSBgDIecdxCg&sessionId=bmZFhfliKMsM). Node degree score of all the genes shown in this network was 4, except *col-169* (node degree: zero)

**Figure 6.**
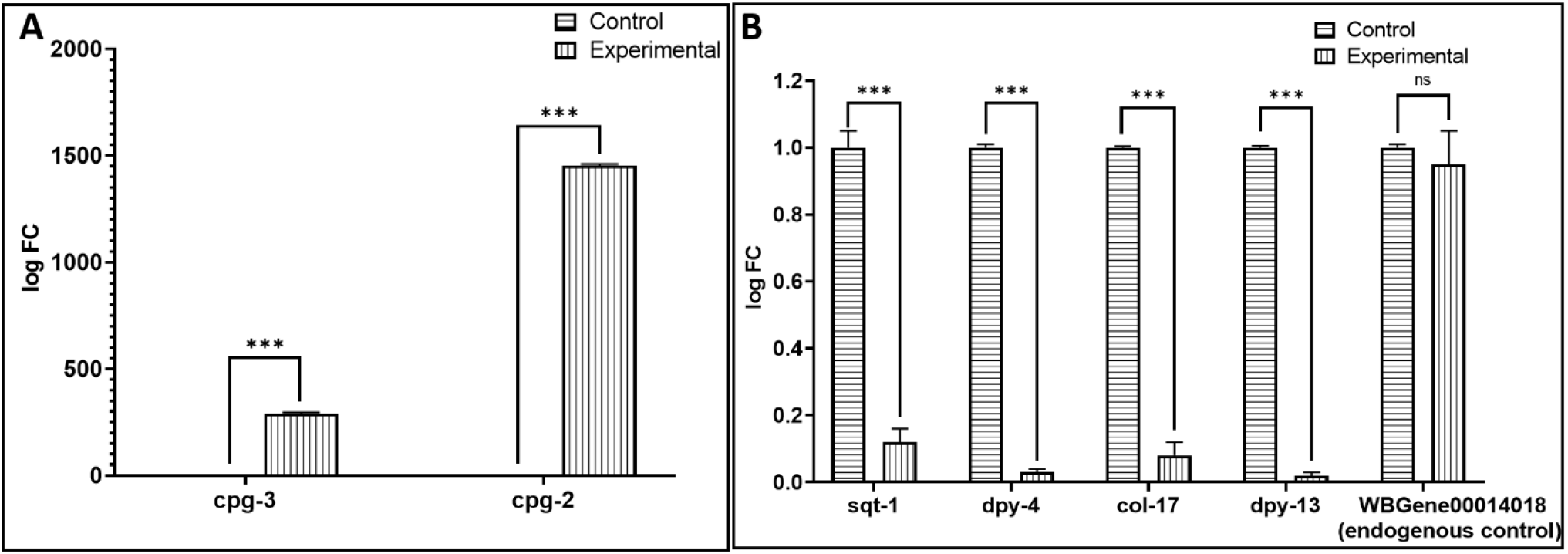
Confirmation of differential expression of selected (A) upregulated and (B) downregulated genes in *W. somnifera-*exposed *C. elegans* through RT-PCR. *** p ≤ 0.001, ns: not significant

**Table 3.** CytoHubba ranking of the top five upregulated high node degree genes.

| No. | Gene ID | Gene symbol | No. of methods ranking this protein among top five | Names of 12 ranking methods of CytoHubba and rank score provided by them |  |  |  |  |  |  |  |  |  |  |  |
| --- | --- | --- | --- | --- | --- | --- | --- | --- | --- | --- | --- | --- | --- | --- | --- |
|  |  |  |  | Degree | MNC | DMNC | MCC | Bottleneck | Eccentricity | Closeness | Radially | Betweenness | Stress | CC | E |
| 1 | WBGene00006929 | *vit-5* | 12 | 8 | 8 | 0.758 | 2880 | 1 | 0.5 | 8.5 | 2.222 | 0.571 | 4 | 0.928 | 3.993 |
| 2 | WBGene00016636 | *perm-2* | 10 | 9 | 9 | - | 2904 | 2 | 1 | 9 | 2.333 | 3.401 | 14 | - | 4.101 |
| 3 | WBGene00011063 | *cpg-3* | 9 | 9 | 9 | - | 2904 | - | 1 | 9 | 2.333 | 3.401 | 14 | - | 4.1 |
| 4 | WBGene00015102 | *cpg-2* | 9 | 8 | 8 | - | - | 1 | 0.5 | 8.5 | 2.222 | 2.452 | 10 | - | 3.982 |
| 5 | WBGene00009035 | F22B3.4 | 8 | 8 | 8 | - | - | 1 | 0.5 | 8.5 | 2.222 | 2.452 | 10 | - | - |

\*- indicates that the given method did not rank that particular gene among top five.

The PPI network for the downregulated genes (Figure 5A) contained 29 nodes connected through 30 edges, with an average node degree of 2.07. Since the number of edges (30) in this PPI network is 30-fold higher than expected (01), this network can be believed to possess significantly more interactions among the member proteins than what can be anticipated for a random set of proteins of same sample size and degree distribution. We arranged members of this PPI network in decreasing order of node degree (Table S4), and those with a score of ≥5 were subjected to ranking by different cytoHubba methods. Then we looked for genes which appeared among the top-6 ranked candidates by ≥6 cytoHubba methods, and six such shortlisted genes (Table 4) were further checked for interactions among themselves followed by cluster analysis (Figure 5B), which showed them to be networked with an average node degree score of 3.88. This network also possessed more edges (10) as against expected (0) for any similar random set of proteins. Four (*sqt-1, dpy-4, dpy-13,* and *col-17*) of these six genes belonged to a single cluster (each with a node degree score of 4), and were chosen for further RT-PCR validation. PCR assay did confirm downregulation of these proteins in extract-exposed worms (Figure 6B).

**Table 4.** CytoHubba ranking of the top six downregulated high node degree genes.

| No. | Gene ID | Gene symbol | No. of methods ranking these proteins among top six | Names of 12 ranking methods of CytoHubba and rank score provided by them |  |  |  |  |  |  |  |  |  |  |  |
| --- | --- | --- | --- | --- | --- | --- | --- | --- | --- | --- | --- | --- | --- | --- | --- |
|  |  |  |  | Degree | MNC | DMNC | MCC | Bottleneck | Eccentricity | Closeness | Radially | Betweenness | Stress | CC | E |
| 1 | WBGene00005016 | *sqt-1* | 12 | 5 | 5 | 0.648 | 120 | 1 | 0.5 | 5 | 0.7 | 0 | 0 | 1 | 3.273 |
| 2 | WBGene00001066 | *dpy-4* | 12 | 5 | 5 | 0.648 | 120 | 1 | 0.5 | 5 | 0.7 | 0 | 0 | 1 | 3.309 |
| 3 | WBGene00000606 | *col-17* | 12 | 5 | 5 | 0.648 | 120 | 1 | 0.5 | 5 | 0.7 | 0 | 0 | 1 | 3.303 |
| 4 | WBGene00000742 | *col-169* | 9 | 5 | 5 | 0.648 | 120 | - | 0.5 | 5 | 0.7 | - | - | 1 | 3.243 |
| 5 | WBGene00001074 | *dpy-13* | 8 | 5 | 5 | 0.648 | 120 | - | 0.5 | 5 | 0.7 | - | - | - | 3.251 |
| 6 | WBGene00000618 | *col-41* | 8 | 5 | 5 | 0.648 | 120 | - | 0.5 | 5 | 0.7 | - | - | - | 3.285 |

‘-’ indicates that the given method did not rank that particular gene among top six.

### 3.5. Identifying the most important DEG based on their homology with human genome

To have an empirical idea about relevance of results of this study in *C. elegans* for humans with respect to *W. somnifera’s* possible beneficial effect, we conducted a cooccurrence analysis of all the 45 DEG in extract-treated *C. elegans* with human genome. This analysis resulted in identification of 16 genes, whose counterpart is present in humans too (Figure 7). Presence of multiple genes in human genome with homology to those expressed differently in extract-exposed worms indicates a good probability of similar effect on humans as observed in *C. elegans* in this study. Among the DEG, *gfat-2* seemed to have the highest similarity with its homologue in humans. The *C. elegans gfat-2* (glutamine-fructose 6-phosphate aminotransferase-2) is 88.3% similar to *gfat-1* (glutamine-fructose 6-phosphate aminotransferase-1) in amino acid sequence, and the latter is considered as a longevity gene (Ruegenberg et al., 2020). gfat-1 is the key enzyme of the hexosamine pathway, and has been mentioned as a regulator of protein quality control and longevity. Increased functionality of *gfat-1* induces endoplasmic reticulum-associated protein degradation and autophagy, and correlatively extends lifespan and ameliorates a wide spectrum of proteinopathies. It can be said that *W. somnifera* root extract promotes health and extend lifespan of *C. elegans* through endogenous modulation of protein quality control. Upregulation of *gfat-1* in our long-lived worms corroborates with the previously published observation that hexosamine pathway metabolites enhance protein quality control and extend life (Denzel et al., 2014). In mammals too, *gfat* is the rate limiting enzyme of the hexosamine pathway. Discussion on important genes listed in Figure-7 other than *gfat-2* has been done in preceding sections.

**Figure 7.**
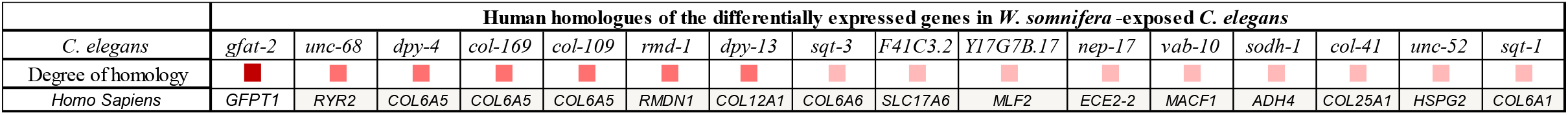
Co-occurrence analysis of differentially expressed genes in extract-treated *C. elegans* with human genome. The darker the shade of the square, the higher the homology between the genes being compared. Genes from left to right are arranged in descending order of homology. Functions of the genes shown here in humans are listed in Table S5.

## 4. Conclusion

This study found the hydroalcoholic root extract of *W. somnifera* (LongeFera^™^) to offer multiple beneficial health effects to the model worm *C. elegans*. Worms incubated with the water-soluble fraction of the extract in absence of any bacterial food displayed more active metabolism and motile behaviour, longer lifespan, and higher fertility. Worm lifespan and healthspan, both were positively influenced by the test extract. Whole transcriptome analysis of the extract-exposed worms revealed the multiple mechanisms through which the extract would have exerted its pro-health effect (Figure 8). Among the major modes of action underlying the observed biological effect of this extract were upregulation of the yolk lipoprotein vitellogenins, remodeling of the apical extracellular matrix, modulation of eggshell permeability, alteration of collagen synthesis, cuticle development and molting cycle. Many of the differentially expressed genes in the extract-fed worms have a homologous counterpart in humans. Considering various parameters like fold-change value, network centrality, cytoHubba ranking, contribution towards more than one function, and homology with human genome, the most important among all DEG are *gfat-2, sqt- 1, dpy-13, dpy-4, cpg-3, vit-5,* and *col-169.* While *W. somnifera* is one of the most popular plants in traditional medicine claimed to have multiple health benefits, molecular mechanisms associated with biological activities of whole extracts from this plant are not that well investigated. Although many natural products have been reported to impart pro-longevity effect on different biological model organisms based on lifespan assays, which is a conventional method to monitor ageing, it should be noted that a compound or formulation that extends lifespan may not necessarily maintain health (Zavagno et al., 2024). The present study has demonstrated *W. somnifera*’s ability to extend lifespan of the model worm *C. elegans* while simultaneously supporting healthy aging, and allowing progeny formation in absence of any bacterial food. Results of this study validate the pro-health efficacy *W. somnifera* claimed in traditional medicine systems like *Ayurved*, and strengthen the candidature of this plant as a potential phytopharmaceutical for healthy ageing.

**Figure 8.**
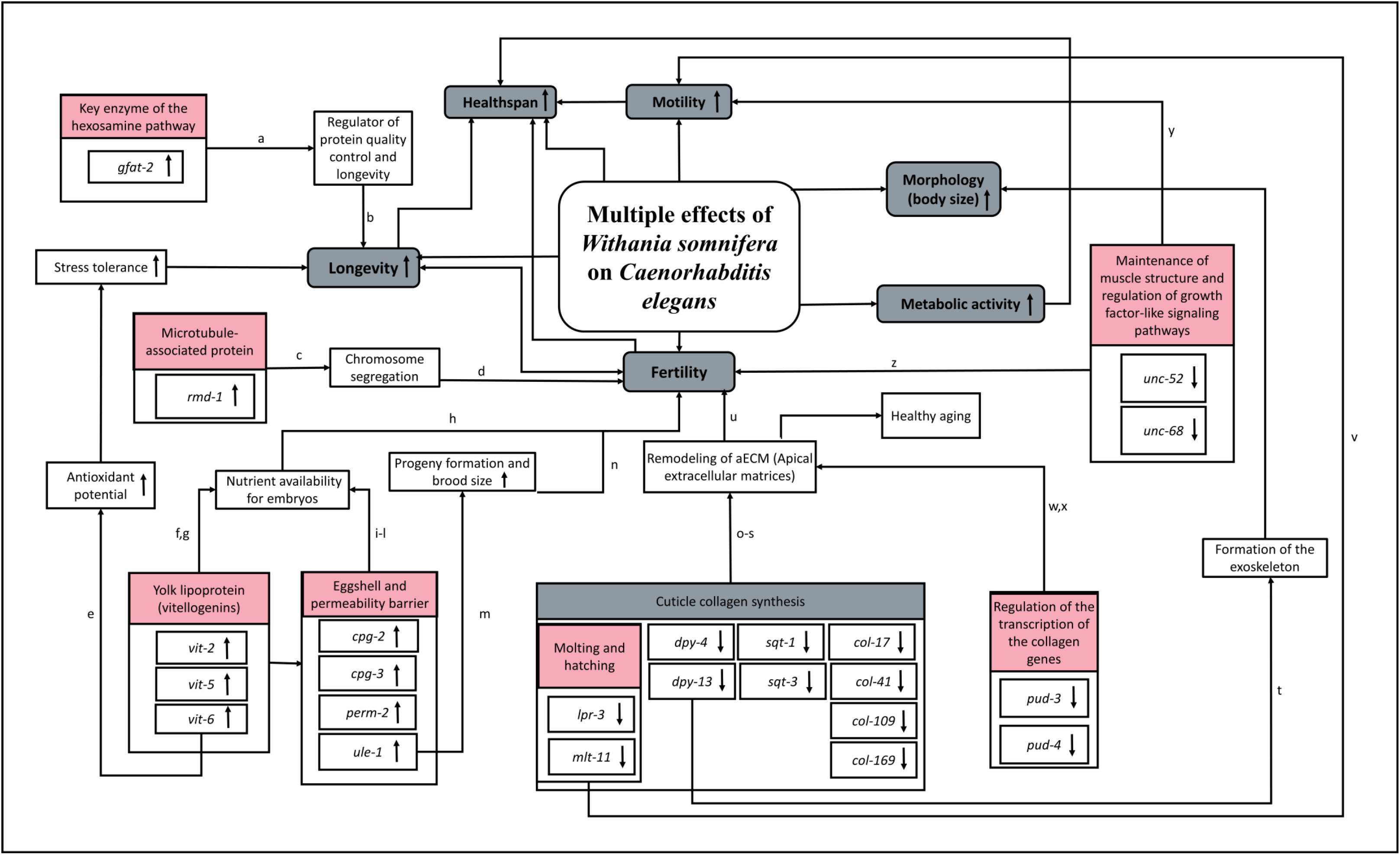
A schematic summary of multiple effects of *W. somnifera* on *C. elegans*. [^a^Ruegenberg et al., 2020; ^b^Denzel et al., 2014; ^c^Oishi et al., 2007; ^d^Juanico et al., 2021; ^e^Nakamura et al., 1999; ^f^Seah et al., 2015; ^g^Perez and Lehner, 2019; ^h^Sornda et al., 2019; ^i^Olson et al., 2012; ^j^Olson et al., 2006; ^k^Kudron et al., 2013; ^l^González et al., 2018; ^m^Maeda et al., 2001; ^n^Green et al., 2011; °Forman-Rubinsky et al., 2017; ^p^Clancy et al., 2023; ^q^Kramer et al., 1988; ^r^Birnbaum et al., 2023; ^s^Nyström et al., 2002; ^t^von Mende et al., 1988; ^u^Adams et al., 2023; ^v^Ragle et al., 2022; ^w^Ding et al., 2013; ^x^Ding et al., 2013; ^y^Merz et al., 2003; ^z^Maryon et al., 1996]

## Supplementary Materials

Table S1. Quantification of extracted RNA, library, and insert size; Table S2. Temperature profile for RT-PCR assay; Table S3. Node degree score of the upregulated genes in *W. somnifera-*exposed *C. elegans*; Table S4. Node degree score of the downregulated genes in *W. somnifera-*exposed *C. elegans*; Table S5. Functions of human genes which are homologous to the differently expressed genes in extract-treated worms; Figure S1. HPLC profile and marker identification in *W. somnifera* root extract; Figure S2. A schematic presentation of the workflow for whole transcriptome and network analysis of WSRE-treated worms; Figure S3. Heat map of DEGs in *W. somnifera*-exposed *C. elegans*.; Figure S4. Volcano Plot of experimental versus control samples; Videos: S1-S4 along with their legend.

## Acknowledgements

We thank NERF (Nirma Education and Research Foundation), Ahmedabad, for infrastructural support, and for providing stipend to NT. GG acknowledges the scholarship from the Government of Gujarat via their SHODH scheme.

## References

1. Abete-Luzi P, Eisenmann DM. Regulation of C. elegans L4 cuticle collagen genes by the heterochronic protein LIN-29. Genesis. 2018;56(5). 10.1002/dvg.23106

2. Adams JR, Pooranachithra M, Jyo EM, Zheng SL, Goncharov A, Crew JR, Kramer JM, Jin Y, Ernst AM, Chisholm AD. Nanoscale patterning of collagens in C. elegans apical extracellular matrix. Nature Communications. 2023;14(1):7506. 10.1038/s41467-023-43058-9

3. Aitlhadj L, Stürzenbaum SR. The use of FUdR can cause prolonged longevity in mutant nematodes. Mechanisms of Ageing and Development. 2010;131(5):364–5. 10.1016/j.mad.2010.03.002

4. Amrit FR, Ratnappan R, Keith SA, Ghazi A. The C. elegans lifespan assay toolkit. Methods. 2014;68(3):465–75. 10.1016/j.ymeth.2014.04.002

5. Anderson SM, Cheesman HK, Peterson ND, Salisbury JE, Soukas AA, Pukkila-Worley R. The fatty acid oleate is required for innate immune activation and pathogen defense in Caenorhabditis elegans. PLoS Pathogens. 2019;15(6):e1007893. 10.1371/journal.ppat.1007893

6. Andrews, S. FastQC: A Quality Control Tool for High Throughput Sequence Data. 2010. Available online: https://www.bioinformatics.babraham.ac.uk/projects/fastqc/

7. Arya U, Das CK, Subramaniam JR. Caenorhabditis elegans for preclinical drug discovery. Current Science. 2010:1669–80.

8. Baker ME. Is vitellogenin an ancestor of apolipoprotein B-100 of human low-density lipoprotein and human lipoprotein lipase?. Biochemical Journal. 1988;255(3):1057–60. 10.1042/bj2551057

9. Baliga MS, Meera S, Shivashankara AR, Palatty PL, Haniadka R. The health benefits of Indian traditional ayurvedic Rasayana (Anti-Aging) drugs: a review. Foods and Dietary Supplements in the Prevention and Treatment of Disease in Older Adults. 2015:151–61. 10.1016/B978-0-12-418680-4.00016-6

10. Basudkar V, Gujrati G, Ajgaonkar S, Gandhi M, Mehta D, Nair S. Emerging vistas for the nutraceutical withania somnifera in inflammaging. Pharmaceuticals. 2024;17(5):597. 10.3390/ph17050597

11. Bhalani DV, Nutan B, Kumar A, Singh Chandel AK. Bioavailability enhancement techniques for poorly aqueous soluble drugs and therapeutics. Biomedicines. 2022;10(9):2055. doi: 10.3390/biomedicines10092055

12. Bharti VK, Malik JK, Gupta RC. Ashwagandha: multiple health benefits. In: Nutraceuticals 2016(pp. 717–733). Academic Press. DOI: 10.1016/B978-0-12-802147-7.00052-8

13. Birnbaum SK, Cohen JD, Belfi A, Murray JI, Adams JR, Chisholm AD, Sundaram MV. The proprotein convertase BLI-4 promotes collagen secretion during assembly of the Caenorhabditis elegans cuticle. bioRxiv. 2023 10.1101/2023.06.06.542650

14. Blighe K, Rana S, Lewis M. EnhancedVolcano: Publication-ready volcano plots with enhanced colouring and labeling. R package version. 2019;1(0):10–8129.

15. Calahorro F, Ruiz-Rubio M. Caenorhabditis elegans as an experimental tool for the study of complex neurological diseases: Parkinson’s disease, Alzheimer’s disease and autism spectrum disorder. Invertebrate Neuroscience. 2011; 11:73–83. 10.1007/s10158-011-0126-1

16. Chen, S.; Zhou, Y.; Chen, Y.; Gu, J. fastp: An ultra-fast all-in-one FASTQ preprocessor. Bioinformatics 2018, 34, i884–90.

17. Chin, C.H.; Chen, S.H.; Wu, H.H.; Ho, C.W.; Ko, M.T.; Lin, C.Y. cytoHubba: Identifying hub objects and sub-networks from complex interactome. BMC System Biology. 2014, 8, S11.

18. Clancy JC, Vo AA, Myles KM, Levenson MT, Ragle JM, Ward JD. Experimental considerations for study of C. elegans lysosomal proteins. G3: Genes, Genomes, Genetics. 2023;13(4):jkad032. 10.1093/g3journal/jkad032

19. Cohen JD, Cadena del Castillo CE, Serra ND, Kaech A, Spang A, Sundaram MV. The Caenorhabditis elegans Patched domain protein PTR-4 is required for proper organization of the precuticular apical extracellular matrix. Genetics. 2021;219(3):iyab132. DOI: 10.1093/genetics/iyab132

20. Corsi AK, Wightman B, Chalfie M. A transparent window into biology: a primer on Caenorhabditis elegans. Genetics. 2015;200(2):387–407. 10.1534/genetics.115.180133

21. Cui Y, McBride SJ, Boyd WA, Alper S, Freedman JH. Toxicogenomic analysis of Caenorhabditis elegans reveals novel genes and pathways involved in the resistance to cadmium toxicity. Genome Biology. 2007; 8(6):R122. 10.1186/gb-2007-8-6-r122

22. Denzel MS, Storm NJ, Gutschmidt A, Baddi R, Hinze Y, Jarosch E, Sommer T, Hoppe T, Antebi A. Hexosamine pathway metabolites enhance protein quality control and prolong life. Cell. 2014;156(6):1167–78. 10.1016/j.cell.2014.01.061

23. Ding YH, Du YG, Luo S, Li YX, Li TM, Yoshina S, Wang X, Klage K, Mitani S, Ye K, Dong MQ. Characterization of PUD-1 and PUD-2, two proteins up-regulated in a long-lived daf-2 mutant. PLoS One. 2013;8(6):e67158. 10.1371/annotation/6b155146-de73-4733-83b0-62224d84717e

24. Dobin A, Davis CA, Schlesinger F, Drenkow J, Zaleski C, Jha S, Batut P, Chaisson M, Gingeras TR. STAR: ultrafast universal RNA-seq aligner. Bioinformatics. 2013;29(1):15–21. 10.1093/bioinformatics/bts635

25. Ewels P, Magnusson M, Lundin S, Käller M. MultiQC: summarize analysis results for multiple tools and samples in a single report. Bioinformatics. 2016;32(19):3047–8. 10.1093/bioinformatics/btw354

26. Forman-Rubinsky R, Cohen JD, Sundaram MV. Lipocalins are required for apical extracellular matrix organization and remodeling in Caenorhabditis elegans. Genetics. 2017;207(2):625–42. 10.1534/genetics.117.300207

27. Ge SX, Jung D, Yao R. ShinyGO: a graphical gene-set enrichment tool for animals and plants. Bioinformatics. 2020;36(8):2628–9. 10.1093/bioinformatics/btz931

28. González DP, Lamb HV, Partida D, Wilson ZT, Harrison MC, Prieto JA, Moresco JJ, Diedrich JK, Yates III JR, Olson SK. CBD-1 organizes two independent complexes required for eggshell vitelline layer formation and egg activation in C. elegans. Developmental Biology. 2018;442(2):288–300. 10.1016/j.ydbio.2018.08.005

29. Green RA, Kao HL, Audhya A, Arur S, Mayers JR, Fridolfsson HN, Schulman M, Schloissnig S, Niessen S, Laband K, Wang S. A high-resolution C. elegans essential gene network based on phenotypic profiling of a complex tissue. Cell. 2011;145(3):470–82. doi: 10.1016/j.cell.2011.03.037

30. Hamid R, Rotshteyn Y, Rabadi L, Parikh R, Bullock P. Comparison of alamar blue and MTT assays for high through-put screening. Toxicology In Vitro. 2004;18(5):703–10. 10.1016/j.tiv.2004.03.012

31. Hardaker L. A., E. Singer, R. Kerr, G. Zhou, and W. R. Schafer, 2001 Serotonin modulates locomotory behavior and coordinates egg-laying and movement in Caenorhabditis elegans. Journal of Neurobiology. 49: 303–313. PMID: 11745666

32. Herndon LA, Schmeissner PJ, Dudaronek JM, Brown PA, Listner KM, Sakano Y, Paupard MC, Hall DH, Driscoll M. Stochastic and genetic factors influence tissue-specific decline in ageing C. elegans. Nature. 2002;419(6909):808–14. 10.1038/nature01135

33. Joshi C, Patel P, Palep H, Kothari V. Validation of the anti-infective potential of a polyherbal ‘Panchvalkal’ preparation, and elucidation of the molecular basis underlining its efficacy against Pseudomonas aeruginosa. BMC Complementary and Alternative Medicine. 2019 (1):1–5. 10.1186/s12906-019-2428-5

34. Juanico IY, Meyer CM, McCarthy JE, Gong T, McNally FJ. Paternal mitochondria from an rmd-2, rmd-3, rmd-6 triple mutant are properly positioned in the C. elegans zygote. Micropublication Biology. 2021;2021. 10.17912%2Fmicropub.biology.000422

35. Kassambara A, Kassambara MA. Package ‘ggpubr’. R package version 0.1. 2020;6(0).

36. Klass MR. Aging in the nematode Caenorhabditis elegans: major biological and environmental factors influencing life span. Mechanisms of Ageing and Development. 1977;6:413–29. 10.1016/0047-6374(77)90043-4

37. Kojima T, Kamei H, Aizu T, Arai Y, Takayama M, Nakazawa S, Ebihara Y, Inagaki H, Masui Y, Gondo Y, Sakaki Y. Association analysis between longevity in the Japanese population and polymorphic variants of genes involved in insulin and insulin-like growth factor 1 signaling pathways. Experimental Gerontology. 2004;39(11-12):1595–8. 10.1016/j.exger.2004.05.007

38. Kramer JM, Johnson JJ, Edgar RS, Basch C, Roberts S. The sqt-1 gene of C. elegans encodes a collagen critical for organismal morphogenesis. Cell. 1988;55(4):555–65. 10.1016/0092-8674(88)90214-

39. Kuboyama T, Tohda C, Komatsu K. Effects of Ashwagandha (roots of Withania somnifera) on neurodegenerative diseases. Biological and Pharmaceutical Bulletin. 2014; 37(6):892–7. 10.1248/bpb.b14-00022

40. Kudron M, Niu W, Lu Z, Wang G, Gerstein M, Snyder M, Reinke V. Tissue-specific direct targets of Caenorhabditis elegans Rb/E2F dictate distinct somatic and germline programs. Genome Biology. 2013;14:1–7.

41. Kumar R, Gupta K, Saharia K, Pradhan D, Subramaniam JR. Withania somnifera root extract extends lifespan of Caenorhabditis elegans. Annals of Neurosciences. 2013;20(1):13. 10.5214/ans.0972.7531.200106

42. Lang AE, Lundquist EA. The Collagens DPY-17 and SQT-3 Direct Anterior–Posterior Migration of the Q Neuroblasts in C. elegans. Journal of Developmental Biology. 2021;9(1):7. 10.3390/jdb9010007

43. Li H, Zhang S. Functions of vitellogenin in eggs. Oocytes: Maternal Information and Functions. 2017:389–401. 10.1007/978-3-319-60855-6_17

44. Liao Y, Smyth GK, Shi W. featureCounts: an efficient general purpose program for assigning sequence reads to genomic features. Bioinformatics. 2014;30(7):923–30. 10.1093/bioinformatics/btt656

45. Longhin EM, El Yamani N, Rundén-Pran E, Dusinska M. The alamar blue assay in the context of safety testing of nanomaterials. Frontiers in Toxicology. 2022;4:981701. 10.3389/ftox.2022.981701

46. Maeda I, Kohara Y, Yamamoto M, Sugimoto A. Large-scale analysis of gene function in Caenorhabditis elegans by high-throughput RNAi. Current Biology. 2001; 11(3):171–6. 10.1016/S0960-9822(01)00052-5

47. Maryon EB, Coronado R, Anderson P. unc-68 encodes a ryanodine receptor involved in regulating C. elegans body-wall muscle contraction. The Journal of Cell Biology. 1996;134(4):885–93.

48. Mehta V, Chander H, Munshi A. Mechanisms of anti-tumor activity of Withania somnifera (Ashwagandha). Nutrition and Cancer. 2021;73(6):914–26. 10.1080/01635581.2020.1778746

49. Merz DC, Alves G, Kawano T, Zheng H, Culotti JG. UNC-52/perlecan affects gonadal leader cell migrations in C. elegans hermaphrodites through alterations in growth factor signaling. Developmental Biology. 2003;256(1):174–87. doi:10.1016/S0012-1606(03)00014-9

50. Mitchell DH, Stiles JW, Santelli J, Sanadi DR. Synchronous growth and aging of Caenorhabditis elegans in the presence of fluorodeoxyuridine. Journal of Gerontology. 1979;34(1):28–36. 10.1093/geronj/34.1.28

51. Mudunuri, U., Che, A., Yi, M., & Stephens, R. M. (2009). bioDBnet: the biological database network. Bioinformatics, 25(4), 555–556.

52. Nagy S., Y.-C. Huang, M. J. Alkema, and D. Biron, 2015 Caenorhabditis elegans exhibit a coupling between the defecation motor program and directed locomotion. Scientific Reports 5: 17174. 10.1038/srep17174. PMID: 26597056

53. Nakamura A, Yasuda K, Adachi H, Sakurai Y, Ishii N, Goto S. Vitellogenin-6 is a major carbonylated protein in aged nematode, Caenorhabditis elegans. Biochemical and Biophysical Research Communications. 1999;264(2):580–3. 10.1006/bbrc.1999.1549

54. Novelli J, Page AP, Hodgkin J. The C terminus of collagen SQT-3 has complex and essential functions in nematode collagen assembly. Genetics. 2006;172(4):2253–67. DOI: 10.1534/genetics.105.053637

55. Nyström J, Shen ZZ, Aili M, Flemming AJ, Leroi A, Tuck S. Increased or decreased levels of Caenorhabditis elegans lon-3, a gene encoding a collagen, cause reciprocal changes in body length. Genetics. 2002;161(1):83–97. 10.1093/genetics/161.1.83

56. Oishi K, Okano H, Sawa H. RMD-1, a novel microtubule-associated protein, functions in chromosome segregation in Caenorhabditis elegans. The Journal of Cell Biology. 2007;179(6):1149–62. http://www.jcb.org/cgi/doi/10.1083/jcb.200705108

57. Olson SK, Bishop JR, Yates JR, Oegema K, Esko JD. Identification of novel chondroitin proteoglycans in Caenorhabditis elegans: embryonic cell division depends on CPG-1 and CPG-2. The Journal of Cell Biology. 2006;173(6):985–94. 10.1083/jcb.200603003

58. Olson SK, Greenan G, Desai A, Müller-Reichert T, Oegema K. Hierarchical assembly of the eggshell and permeability barrier in C. elegans. Journal of Cell Biology. 2012;198(4):731–48.

59. Patel P, Joshi C, Kothari V. Anti-pathogenic efficacy and molecular targets of a polyherbal wound-care formulation (Herboheal) against Staphylococcus aureus. Infectious Disorders-Drug Targets (Formerly Current Drug Targets-Infectious Disorders). 2019;19(2):193–206. 10.2174/1871526518666181022112552

60. Perez MF, Lehner B. Vitellogenins-yolk gene function and regulation in Caenorhabditis elegans. Frontiers in Physiology. 2019;10:471381. 10.3389/fphys.2019.01067

61. Peterson ND, Cheesman HK, Liu P, Anderson SM, Foster KJ, Chhaya R, Perrat P, Thekkiniath J, Yang Q, Haynes CM, Pukkila-Worley R. The nuclear hormone receptor NHR-86 controls anti-pathogen responses in C. elegans. PLoS Genetics. 2019;15(1):e1007935. 10.1371/journal.pgen.1007935

62. Pickett CL, Dietrich N, Chen J, Xiong C, Kornfeld K. Mated progeny production is a biomarker of aging in Caenorhabditis elegans. G3. 2013; 3(12):2219–32. doi: 10.1534/g3.113.008664

63. Ragle JM, Levenson MT, Clancy JC, Vo AA, Pham V, Ward JD. The conserved, secreted protease inhibitor MLT-11 is necessary for C. elegans molting and embryogenesis. bioRxiv. 2022:2022–06. 10.1101/2022.06.29.498124

64. Ravi B, Garcia J, Collins KM. The HSN egg-laying command neurons regulate the defecation motor program in Caenorhabditis elegans: Integration. MicroPublication Biology. 2019;2019. 10.17912%2Fmicropub.biology.000095

65. Robinson, M. D., McCarthy, D. J., & Smyth, G. K. (2010). edgeR: a Bioconductor package for differential expression analysis of digital gene expression data. Bioinformatics, 26(1), 139–140.

66. Rogalski TM, Mullen GP, Bush JA, Gilchrist EJ, Moerman DG. UNC-52/perlecan isoform diversity and function in Caenorhabditis elegans. Biochemical Society Transactions. 2001;29(2):171–6.

67. Ruegenberg S, Horn M, Pichlo C, Allmeroth K, Baumann U, Denzel MS. Loss of GFAT-1 feedback regulation activates the hexosamine pathway that modulates protein homeostasis. Nature Communications. 2020;11(1):687. 10.1038/s41467-020-14524-5

68. Sandhu A, Badal D, Sheokand R, Tyagi S, Singh V. Specific collagens maintain the cuticle permeability barrier in Caenorhabditis elegans. Genetics. 2021;217(3):iyaa047. 10.1093/genetics/iyaa047

69. Seah NE, de Magalhaes Filho CD, Petrashen AP, Henderson HR, Laguer J, Gonzalez J, Dillin A, Hansen M, Lapierre LR. Autophagy-mediated longevity is modulated by lipoprotein biogenesis. Autophagy. 2016;12(2):261–72. 10.1080/15548627.2015.1127464

70. Sengupta P, Agarwal A, Pogrebetskaya M, Roychoudhury S, Durairajanayagam D, Henkel R. Role of Withania somnifera (Ashwagandha) in the management of male infertility. Reproductive Biomedicine Online. 2018;36(3):311–26. 10.1016/j.rbmo.2017.11.007

71. Shannon, P.; Markiel, A.; Ozier, O.; Baliga, N.S.; Wang, J.T.; Ramage, D.; Amin, N.; Schwikowski, B.; Ideker, T. Cytoscape: A software environment for integrated models of biomolecular interaction networks. Genome Research. 2003, 13, 2498–2504.

72. Sharma PK, Kumar L, Goswami Y, Pujani M, Dikshit M, Tandon R. The aqueous root extract of Withania somnifera ameliorates LPS-induced inflammatory changes in the in vitro cell-based and mice models of inflammation. Frontiers in Pharmacology. 2023;14:1139654. 10.3389/fphar.2023.1139654

73. Sifri CD, Begun J, Ausubel FM. The worm has turned–microbial virulence modeled in Caenorhabditis elegans. Trends in Microbiology. 2005;13(3):119–27. 10.1016/j.tim.2005.01.003

74. Singh M, Jayant K, Singh D, Bhutani S, Poddar NK, Chaudhary AA, Khan SU, Adnan M, Siddiqui AJ, Hassan MI, Khan FI. Withania somnifera (L.) Dunal (Ashwagandha) for the possible therapeutics and clinical management of SARS-CoV-2 infection: Plant-based drug discovery and targeted therapy. Frontiers in Cellular and Infection Microbiology. 2022;12:933824. 10.3389/fcimb.2022.933824

75. Sornda T, Ezcurra M, Kern C, Galimov ER, Au C, de la Guardia Y, Gems D. Production of YP170 vitellogenins promotes intestinal senescence in Caenorhabditis elegans. The Journals of Gerontology: Series A. 2019;74(8):1180–8. 10.1093/gerona/glz067

76. Speers AB, Cabey KA, Soumyanath A, Wright KM. Effects of Withania somnifera (Ashwagandha) on stress and the stress-related neuropsychiatric disorders anxiety, depression, and insomnia. Current Neuropharmacology. 2021;19(9):1468. doi:10.2174/1570159X19666210712151556

77. Suh Y, Atzmon G, Cho MO, Hwang D, Liu B, Leahy DJ, Barzilai N, Cohen P. Functionally significant insulin-like growth factor I receptor mutations in centenarians. Proceedings of the National Academy of Sciences. 2008;105(9):3438–42.

78. Szklarczyk, D.; Gable, A.L.; Lyon, D.; Junge, A.; Wyder, S.; Huerta-Cepas, J.; Simonovic, M.; Doncheva, N.T.; Morris, J.H.; Bork, P.;, et al. STRING v11: Protein–protein association networks with increased coverage, supporting functional discovery in genome-wide experimental datasets. Nucleic Acids Research 2019, 47, D607–13.

79. Tao J, Wu QY, Ma YC, Chen YL, Zou CG. Antioxidant response is a protective mechanism against nutrient deprivation in C. elegans. Scientific Reports. 2017;7(1):43547. DOI: 10.1038/srep43547

80. Tritten L, Braissant O, Keiser J. Comparison of novel and existing tools for studying drug sensitivity against the hookworm Ancylostoma ceylanicum in vitro. Parasitology. 2012;139(3):348–57. doi:10.1017/S0031182011001934

81. United States Pharmacopeial Convention, Ashwagandha root; powdered ashwagandha root; and powdered ashwagandha root extract, U.S. Pharmacopeia, N. Formulary (Eds.), Dietary supplements compendium. Rockville, MD, 2019.

82. Untergasser, A.; Nijveen, H.; Rao, X.; Bisseling, T.; Geurts, R.; Leunissen, J.A. Primer3Plus, an enhanced web interface to Primer3. Nucleic Acids Research. 2007, 35 (Suppl. S2), W71–W74.

83. Vaidya VG, Naik NN, Ganu G, Parmar V, Jagtap S, Saste G, Bhatt A, Mulay V, Girme A, Modi SJ, Hingorani L. Clinical pharmacokinetic evaluation of Withania somnifera (L.) Dunal root extract in healthy human volunteers: A non-randomized, single dose study utilizing UHPLC-MS/MS analysis. Journal of Ethnopharmacology. 2024; 322:117603. 10.1016/j.jep.2023.117603

84. von Mende N, Bird DM, Albert PS, Riddle DL. dpy-13: a nematode collagen gene that affects body shape. Cell. 1988;55(4):567–76. 10.1016/0092-8674(88)90215-2

85. Xu S, Hsiao TI, Chisholm AD. The wounded worm: Using C. elegans to understand the molecular basis of skin wound healing. In: Worm 2012 (Vol. 1, No. 2, pp. 134–138). Taylor & Francis. 10.4161/worm.19501

86. Zavagno G, Raimundo A, Kirby A, Saunter C, Weinkove D. Rapid measurement of ageing by automated monitoring of movement of C. elegans populations. Geroscience. 2024;46(2):2281–93. 10.1007/s11357-023-00998-w

87. Zavagno G, Raimundo A, Kirby A, Saunter C, Weinkove D. Rapid measurement of ageing by automated monitoring of movement of C. elegans populations. Geroscience. 2024;46(2):2281–93. 10.1007/s11357-023-00998-w

88. Zhu G, Yin F, Wang L, Wei W, Jiang L, Qin J. Modeling type 2 diabetes-like hyperglycemia in C. elegans on a microdevice. Integrative Biology. 2016;8(1):30–8. 10.1039/c5ib00243e

89. Zimmerman SM, Hinkson IV, Elias JE, Kim SK. Reproductive aging drives protein accumulation in the uterus and limits lifespan in C. elegans. PLoS Genetics. 2015;11(12):e1005725.

