## Supplementary material for "*Withania somnifera* root extract (LongeFera™) confers beneficial effects on health and lifespan of the model worm *Caenorhabditis elegans*": Legend for Supplementary Videos_W. somnifera_C. elegans.pdf

### Legend to Supplementary Videos

for


We investigated the effect of the hydroalcoholic extract of *Withania somnifera* root on lifespan and healthspan of the model worm *Caenorhabditis elegans*. Positive effect of the extract on worm's lifespan, motility, and fertility can be visualized in the supplementary videos listed below:

**Video S1**: (control well) A worm in M9 buffer (devoid of plant extract) captured at 7$^{th}$ day. S1a and S1b are two different worms from two different wells.

**Video S2**: Worm population in extract (600 μg/mL)-supplemented media, captured on the 7$^{th}$ day. Actively moving progenies are visible (which were absent in corresponding control video 'A'). Adult worms too can be seen exhibiting movement better than their counterparts in 'control' wells.

Comparative assessment of video A vs. B reveals abnormal body movement and smaller body size in control worms, as against bigger body size and normal movement in extract-fed worms.

**Video S3**: Control worm population at 12$^{th}$ day, whereby ~100% worms were dead, as evident from lack of movement. Morphological damage can also be seen.

**Video S4**: Extract-fed worms captured on the 12$^{th}$ day. Actively moving adult as well as progeny worms can be seen. Morphology of the parent worms (i.e. those with whom experiment started) seems normal.

All these videos were captured at the end of the 7$^{th}$ or 12$^{th}$ day of incubation, i.e., the days on which the control population exhibited ~50% and ~100% death, respectively. Videos were captured using a Magnus Camera (5.1 MP) attached to a Labomed Vision 2000 (halogen light source) binocular microscope (4X objective).
