## Supplementary material for "*Withania somnifera* root extract (LongeFera™) confers beneficial effects on health and lifespan of the model worm *Caenorhabditis elegans*": Supplementary File_Withania somnifera_C. elegans_01.08.2024.pdf

### Table S1. Quantification of extracted RNA, library, and insert size

| Sr. no. | Sample | Quantification of extracted RNA | | | Library quantification and insert size analysis | |
|---|---|---|---|---|---|---|
| | | $OD_{260}/OD_{280}$ | $OD_{260}/OD_{230}$ | RIN value | ng/µL | Insert size (bp) |
| 1 | Control | 1.9 | 2 | 8.9 | 39.4 | 321 |
| 2 | Experimental | 1.8 | 0 | 9.6 | 35.6 | 314 |

### Table S2. Temperature profile for RT-PCR assay

| Temperature (°C) | Time (s) | Remarks |
|---|---|---|
| PCR cycles (45 cycles) | | |
| 95 | 15 | Denaturation temperature |
| 59 | 60 | Annealing temperature |
| Melt curve stage | | |
| 95 | 15 | |
| 60 | 60 | |
| 95 | 15 | |

**Table S3. Node degree score of the upregulated genes in *W. somnifera*-exposed *C. elegans***

| Sr. No. | Gene ID/ symbol | Node Degree |
|---|---|---|
| 1 | *cpg-3* | 10 |
| 2 | F22B3.4 | 9 |
| 3 | *cpg-2* | 9 |
| 4 | *perm-2* | 9 |
| 5 | *vit-2* | 8 |
| 6 | *vit-5* | 8 |
| 7 | *vit-6* | 8 |
| 8 | W03F11.1 | 7 |
| 9 | Y62H9A.3 | 7 |
| 10 | *clec-87* | 5 |
| 11 | *rmd-1* | 4 |
| 12 | F15E11.12 | 1 |
| 13 | F15E11.15 | 1 |

Rest 2 genes with node degree score ‘zero’ are not listed.

**Table S4. Node degree score of the downregulated genes in *W. somnifera*-exposed *C. elegans***

| Sr. No. | Gene ID/ symbol | Node Degree |
|---|---|---|
| 1 | *sqt-1* | 8 |
| 2 | *unc-52* | 8 |
| 3 | *col-169* | 6 |
| 4 | *dpy-13* | 6 |
| 5 | *dpy-4* | 6 |
| 6 | *let-805* | 6 |
| 7 | *mlt-11* | 6 |
| 8 | *ttn-1* | 6 |
| 9 | *col-17* | 5 |
| 10 | *col-41* | 5 |
| 11 | *ketn-1* | 5 |
| 12 | *lpr-3* | 5 |
| 13 | *col-109* | 4 |
| 14 | *dig-1* | 4 |
| 15 | *fbn-1* | 4 |
| 16 | *sqt-3* | 4 |
| 17 | *unc-68* | 4 |
| 18 | *vab-10* | 4 |
| 19 | H03E18.1 | 3 |
| 20 | Y65B4BL.1 | 2 |
| 21 | *cdh-12* | 2 |
| 22 | *nep-17* | 2 |
| 23 | F28B4.3 | 1 |

Rest 6 genes with node degree score ‘zero’ are not listed.

**Table S5. Functions of human genes which are homologous to the differently expressed genes in extract-treated worms**

| Sr. No. | Gene symbol | Function |
|---|---|---|
| 1 | GFPT1 | Glutamine--fructose-6-phosphate aminotransferase [isomerizing] 1; Controls the flux of glucose into the hexosamine pathway. Most likely involved in regulating the availability of precursors for N- and O-linked glycosylation of proteins. Regulates the circadian expression of clock genes ARNTL/BMAL1 and CRY1. |
| 2 | RYR2 | Ryanodine receptor 2; Calcium channel that mediates the release of $Ca^{2+}$ from the sarcoplasmic reticulum into the cytosol and thereby plays a key role in triggering cardiac muscle contraction. |
| 3 | COL6A5 | Collagen alpha-5(VI) chain; Collagen VI acts as a cell-binding protein. |
| 4 | COL6A5 | Collagen alpha-5(VI) chain; Collagen VI acts as a cell-binding protein. |
| 5 | COL6A5 | Collagen alpha-5(VI) chain; Collagen VI acts as a cell-binding protein. |
| 6 | RMDN1 | Regulator of microtubule dynamics 1. |
| 7 | COL12A1 | Collagen alpha-1(XII) chain; Type XII collagen interacts with type I collagen-containing fibrils, the COL1 domain could be associated with the surface of the fibrils, and the COL2 and NC3 domains may be localized in the perifibrillar matrix. |
| 8 | COL6A6 | Collagen alpha-6(VI) chain; Collagen VI acts as a cell-binding protein. |
| 9 | SLC17A6 | Vesicular glutamate transporter 2; Mediates the uptake of glutamate into synaptic vesicles at presynaptic nerve terminals of excitatory neural cells. May also mediate the transport of inorganic phosphate. Belongs to the major facilitator superfamily. Sodium/anion cotransporter family. VGLUT subfamily. |
| 10 | MLF2 | Myeloid leukemia factor 2. |
| 11 | ECE2-2 | EEF1AKMT4-ECE2 read through transcript protein; Converts big endothelin-1 to endothelin-1. May also have methyltransferase activity (By similarity). May play a role in amyloid- beta processing (By similarity); In the C-terminal section; belongs to the peptidase M13 family. |
| 12 | MACF1 | Microtubule-actin cross-linking factor 1, isoforms 1/2/3/5; [Isoform 2]: F-actin-binding protein which plays a role in cross-linking actin to other cytoskeletal proteins and also binds to microtubules. |
| 13 | ADH4 | All-trans-retinol dehydrogenase [$NAD^{(+)}$] ADH4; Catalyzes the NAD-dependent oxidation of either all-trans- retinol or 9-cis-retinol. |
| 14 | COL25A1 | Collagen-like Alzheimer amyloid plaque component; Inhibits fibrillization of amyloid-beta peptide during the elongation phase. Has also been shown to assemble amyloid fibrils into protease-resistant aggregates. Binds heparin. |
| 15 | HSPG2 | Heparan sulfate proteoglycan 2. |
| 16 | COL6A1 | Collagen alpha-1(VI) chain; Collagen VI acts as a cell-binding protein. |

(Source: https://version-12-0.string-db.org/cgi/cooccurrence?networkId=bBZLpbKyQ4Xt)

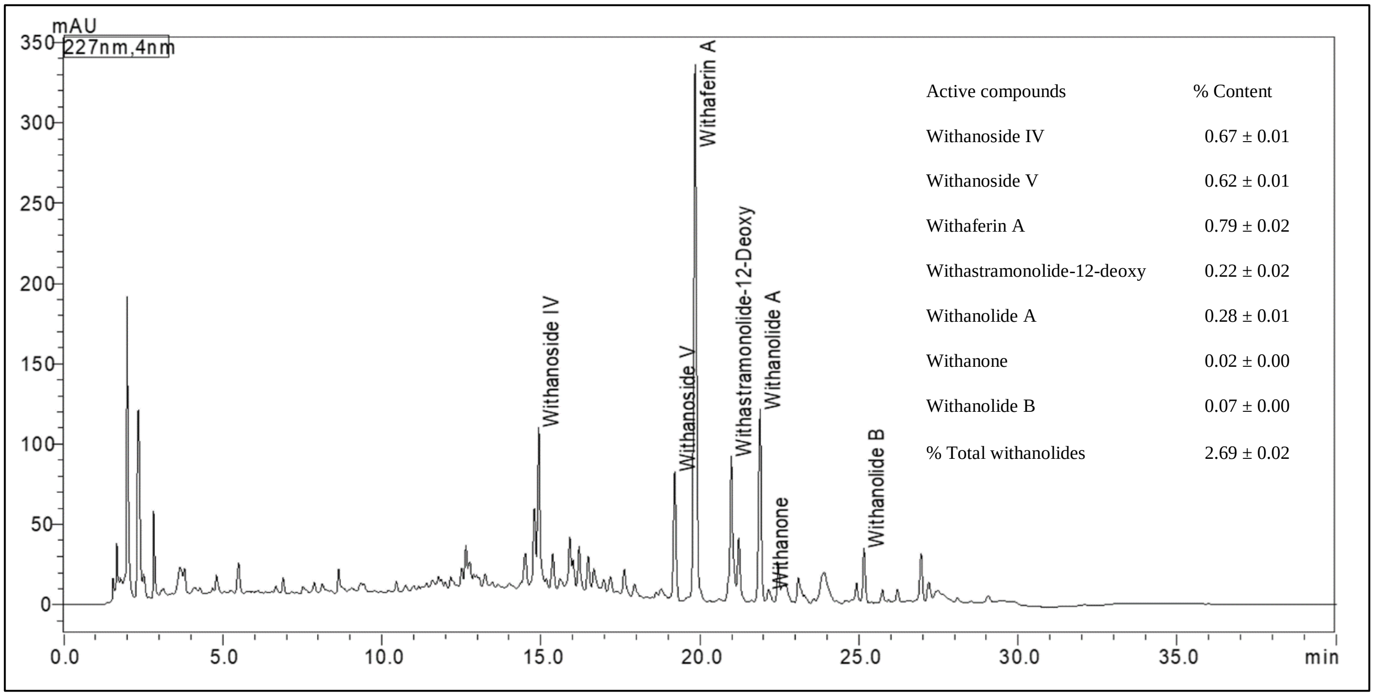


**Figure S1. HPLC profile and marker identification in *W. somnifera* root extract**

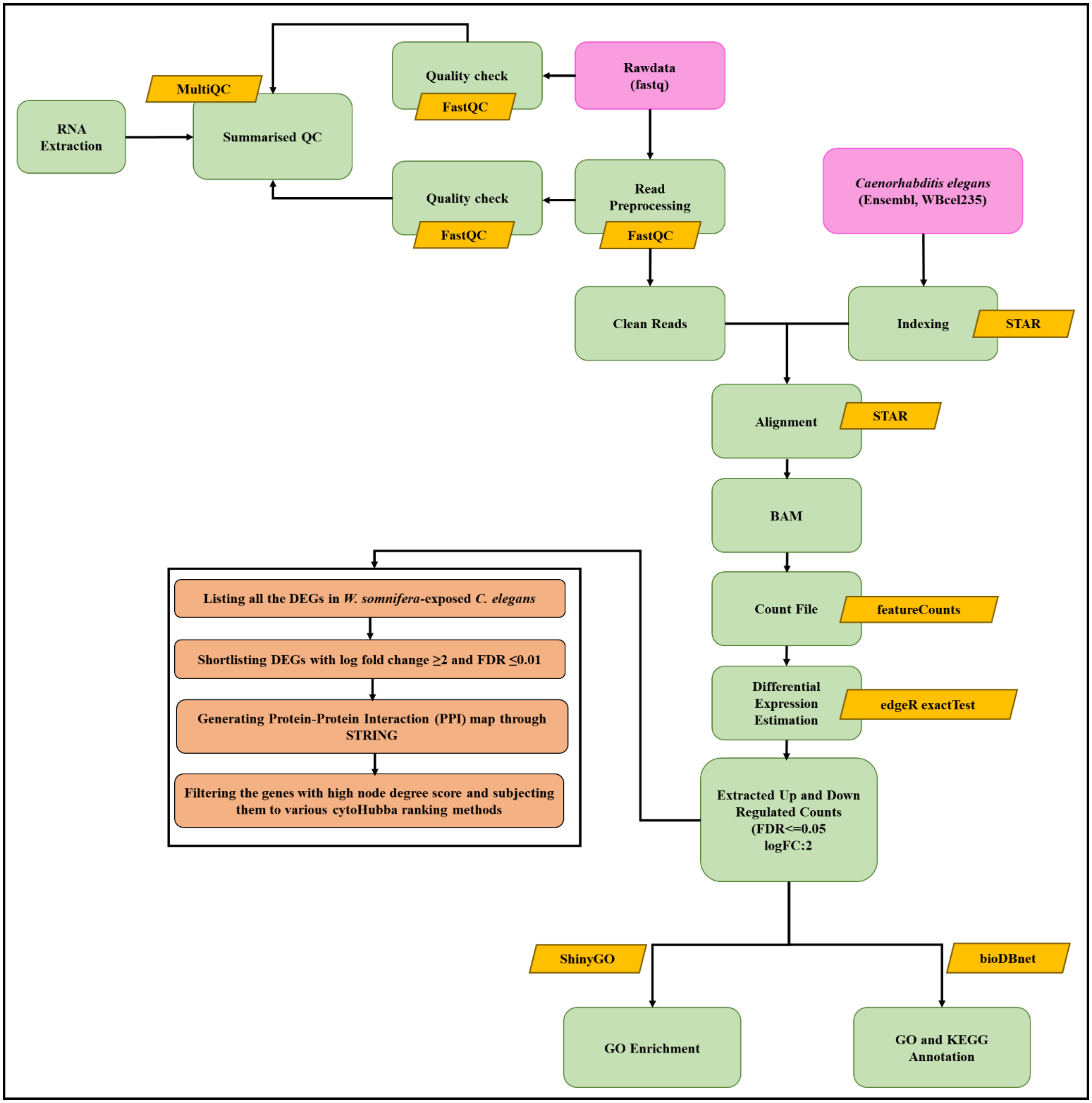


**Figure S2. A schematic presentation of the workflow for whole transcriptome and network analysis of extract-treated worms**

DEG: Differentially Expressed Genes; FDR: False Discovery Rate; log FC: log Fold Change; GO: Gene Ontology (GO); KEGG: Kyoto Encyclopedia of Genes and Genomes

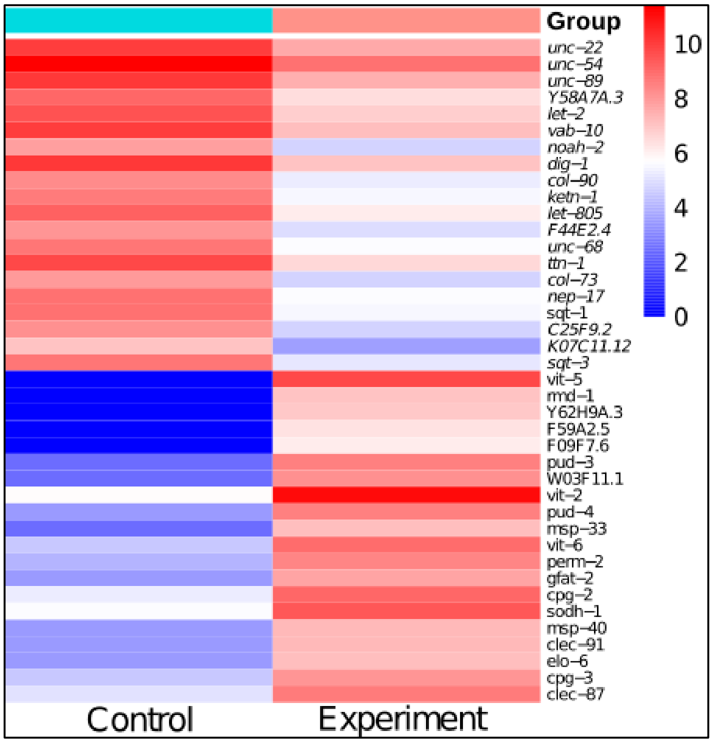


**Figure S3. Heat map of DEGs in *W. somnifera*-exposed *C. elegans*.** Heat map generated using the online software tool ClustVis (https://biit.cs.ut.ee/clust vis/) showing up-regulated and down-regulated genes with FDR<0.01 and log fold change '± 2'.

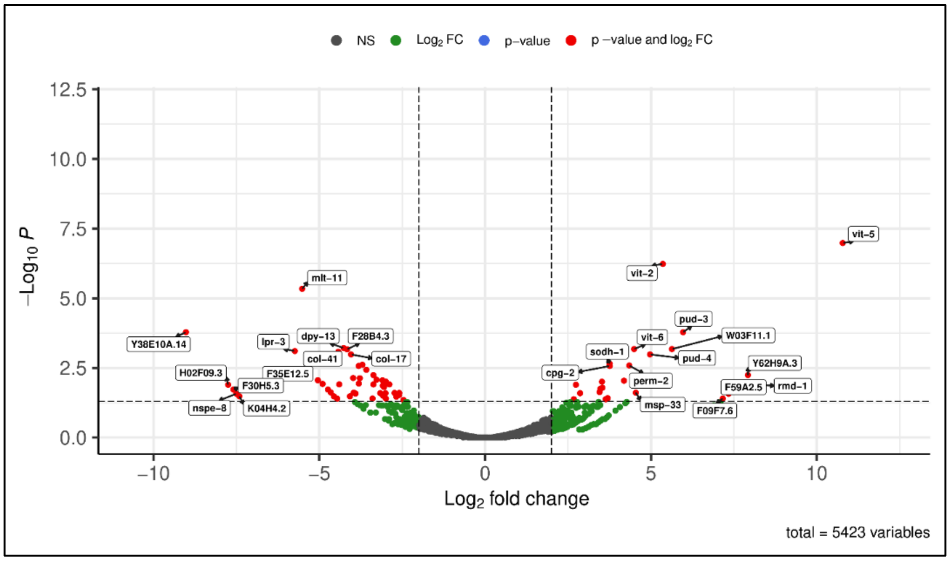


**Figure S4. Volcano Plot of experimental versus control samples.** Volcano plot of expressed genes of experimental culture compared to control culture. The y- axis illustrates −log 10 *p* values, and the x-axis corresponds to a log 2-fold change of gene expression between both cultures. The red points represent differently expressed genes satisfying the dual criteria of FDR≤0.01 and log fold change ≥2.
